# Activation-dependent lentiviruses enable antigen-specific T cell expansion and transduction

**DOI:** 10.64898/2026.05.11.724165

**Authors:** Blake E. Smith, Lindsey M. Draper, Andrea Garmilla, Caleb R. Perez, Nishant Singh, Lucia T. Padilla, Ellen J. K. Xu, Stephanie A. Gaglione, Jiao Shen, Winiffer D. Conce Alberto, Qingyang Henry Zhao, Connor S. Dobson, Kole T. Roybal, Michael Dougan, Michael E. Birnbaum, Stephanie K. Dougan

## Abstract

Cancer immunotherapies rely on tumor-specific T cells, which arise endogenously in most patients with cancer, but can be low frequency and poorly functional. Methods to specifically identify, expand, and manipulate tumor-specific T cells at the rare frequencies found in peripheral blood would enable new immunotherapeutic strategies. Here, we demonstrate an approach to virally transduce polyclonal tumor-reactive T cells across any MHC haplotype and in the absence of knowing the cognate antigen. By generating lentiviral vectors that selectively transduce cells expressing 4-1BB (CD137), a marker of T cell activation, we can transduce antigen-specific T cells with user-defined genetic cargoes that can selectively expand and track individual clonotypes via single-cell sequencing. Anti-4-1BB lentiviruses (4-1BB LVs) encoding therapeutic cargoes can also enhance antigen-specific T cells to extend survival in a xenograft model of human melanoma and transduce tumor-infiltrating T cells from patients with ovarian cancer. Overall, the 4-1BB LV platform targets antigen-specific T cells in a manner agnostic to both the antigen and presenting MHC, with potential applications in adoptive cell therapy manufacturing and TCR identification.

**One Sentence Summary:** Engineered lentiviral vectors targeting 4-1BB selectively activate, expand, and transduce antigen-specific T cells with immunomodulatory cargo.

## INTRODUCTION

Modern cancer immunotherapies harness a patient’s endogenously generated antitumor immune response to achieve tumor control (*1*). Modalities as diverse as immune checkpoint blockade (*2*), cytokine therapy (*3*), and tumor-infiltrating lymphocyte (TIL) therapy (*4*) function by amplifying or reinvigorating tumor-specific T cells. Although these strategies have produced durable responses in patients, their broader impact is constrained by limitations in efficacy and safety (*5*), widespread applicability across tumor types, as well as significant manufacturability challenges, especially for TIL-based therapies (*6*).

Strategies to target existing tumor antigen-specific T cells across a range of rare precursor frequencies would be highly desirable. Tumor-reactive T cells are most abundant in tumor infiltrates (TIL), as evidenced by the clinical success of *ex vivo* expanded TIL therapies (*7-11*). However, tumor-specific T cells also exist in peripheral blood, and their systemic expansion is correlated with response to checkpoint blockade immunotherapies (*12-14*). Tumor-reactive T cells in peripheral blood are more functional than their intratumoral counterparts (*15, 16*), although their rarity has thus far been an impediment to exploiting blood as a source of endogenous T cells (*17, 18*). To date, TIL therapies rely on laborious manufacturing pipelines to generate patient-specific T cells from heavily exhausted T cells present within tumor fragments, and do so without knowledge of *bona fide* tumor reactivity (*4*). Indeed, only a minority of TIL are truly tumor-reactive, with the majority of T cells representing non-tumor-reactive bystander cells that can dominate *ex vivo* expansions with current manufacturing protocols (*19-22*).

Approaches that broadly target T cells would indiscriminately deliver therapeutic cargoes to such bystander T cells as readily as to antigen-specific cells, leading to risks of autoimmunity and reduced efficacy. However, the diverse nature of polyclonal T cell responses makes it infeasible to know the full antigen reactivity of anti-tumor T cells for any given patient. PD-1 blockade tends to target T cells that recognize patient-specific neoantigens derived from mutational processes in the tumor but responses are heterogenous across patients (*23, 24*). Furthermore, the diversity of the MHC locus complicates identification of antigenic epitopes even in patients with shared overexpressed antigens and the same cancer type (*25*). Thus, strategies are needed which can target tumor-reactive T cells in a manner that is agnostic to both tumor antigens and the patient-specific presenting MHC molecules. One strategy to achieve tumor-specificity would be to deliver a CAR or tumor-specific TCR construct to T cells, thereby converting any transduced T cell into a tumor-specific T cell. However, this approach is not feasible for personalized neoantigens, many of which are not cell surface expressed proteins, and the targeting of single antigens can lead to antigen loss as a major mechanism of resistance (*26*). As an alternative approach, we envision delivering immunomodulatory cargoes directly to endogenously generated polyclonal tumor-specific T cells, expanding rare clonotypes in the process, and capturing the immune system’s ability to target multiple antigens in a coordinated CD4 and CD8 T cell response.

Here, we demonstrate methodology to expand and transduce tumor-reactive T cells from polyclonal starting populations absent knowledge of the antigen itself. We coupled an engineered viral fusogen with an agonistic single-chain variable fragment (scFv) that targets 4-1BB (CD137). 4-1BB is a potent costimulatory receptor that promotes T cell survival, proliferation, and cytokine secretion (*27*). 4-1BB is expressed at low levels in resting T cells and becomes transiently upregulated in recently-activated T cells thereby offering a temporal window of selectivity after antigen exposure, agnostic to the identity of the peptide-MHC trigger (*28, 29*). We demonstrate that 4-1BB-targeting LV particles (herein described as “4-1BB LV”) selectively activate and expand antigen-specific clonotypes from very rare precursor frequencies while avoiding off-target T cells. 4-1BB LVs can be used to transfer genetic cargoes that enhance antigen-specific T cell cytotoxicity *in vitro* and *in vivo*. 4-1BB LVs can transduce and expand even dysfunctional T cells as evidenced by efficient gene delivery to T cells isolated from ascites and tumor from patients with ovarian cancer.

## RESULTS

### Developing 4-1BB-targeted lentiviral vectors

Co-display of user-defined ligands and a viral fusogen on the surface of pseudotyped lentiviruses can redirect their tropism relative to lentiviruses expressing the pan-tropic vesicular stomatitis virus glycoprotein (VSVG) alone (*30, 31*). We and others have shown that using a receptor-blinded version of VSVG (“VSVGmut”) as the fusogen expressed alongside a cell-targeting ligand results in lentiviral particles endowed with newfound tropism, efficient infection, and significant selectivity (*30, 32, 33*). We hypothesized that lentiviruses expressing VSVGmut and the endogenous ligand of 4-1BB (4-1BBL, CD137L) or an agonistic single-chain variable fragment (scFv) could achieve selective, 4-1BB-dependent transduction of T cells **(Fig. 1A)**.

**Fig. 1.**
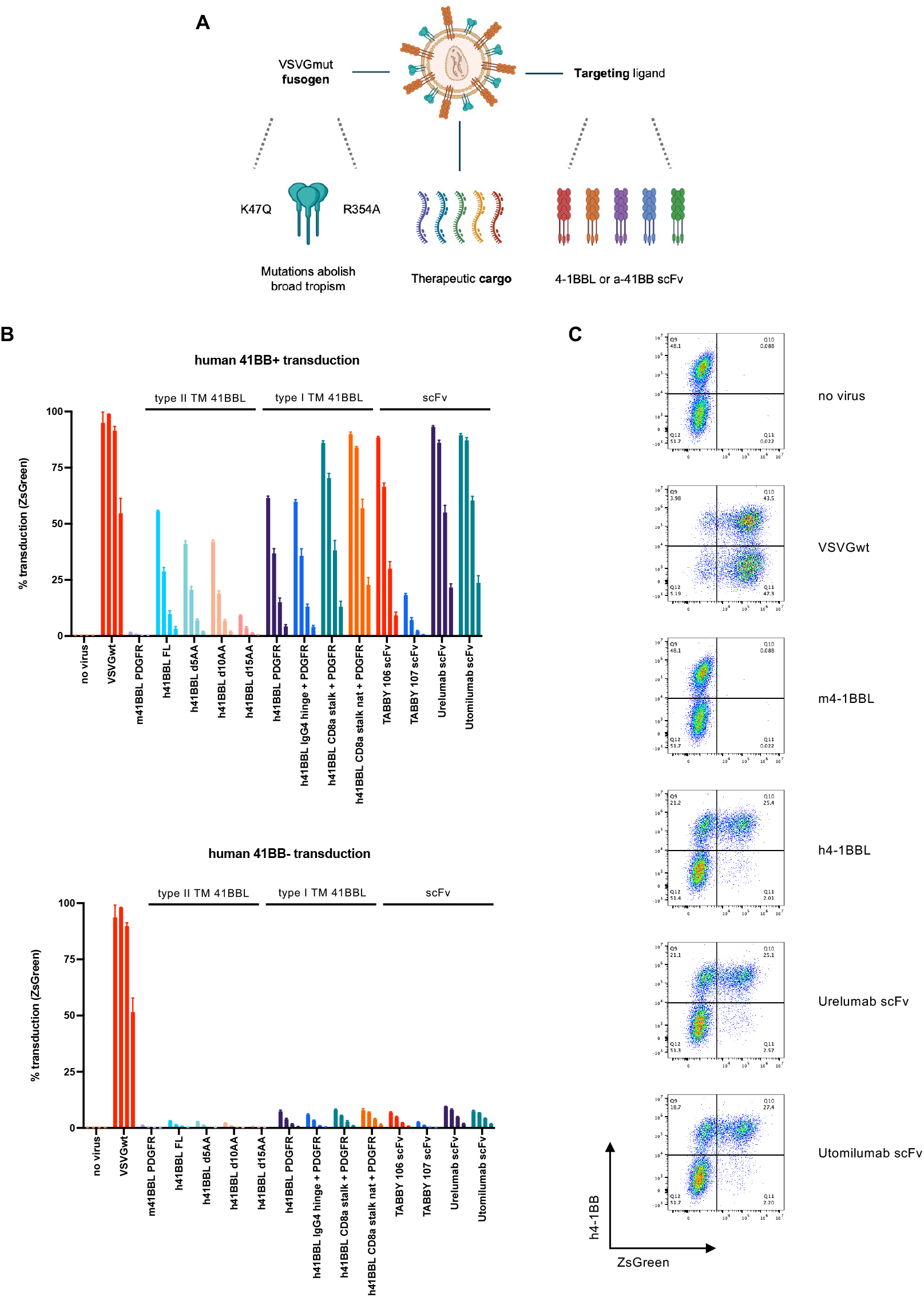
Lentiviral vectors displaying 4-1BBL or anti-4-1BB scFvs selectively target human 4-1BB-expressing Jurkat T cells. **(A)** Pseudotyped lentiviral (LV) vectors designed to target 4-1BB. Co-display of a targeting ligand (e.g. 4-1BBL or anti-4-1BB scFv) alongside VSVGmut (K47Q + R354A) on the surface of an enveloped LV particle can enable cell-specific infection. Desired therapeutic cargo can be packaged inside LV vectors to achieve cell-specific gene delivery. **(B)** Transduction (% ZsGreen+) observed in h4-1BB+ (EGFR+) Jurkat T cells (top) or h4-1BB-(EGFR-) Jurkat T cells (bottom) by construct. Each bar represents triplicate data points per dose of virus added. Each virus was added in a dilution series with dextran, with each bar from left to right (per construct) representing a virus dose of: 1, 0.25, 0.0625, and 0.015625 µL. **(C)** Representative flow plots demonstrating transduction (ZsGreen+) observed in h4-1BB (EGFR+) Jurkat T cells with selected lead candidate constructs and pertinent controls at a single virus dose (0.0625 µL).

Empiric differences in targeting ligands can have drastic effects on the outcome of transduction via the cognate receptor (*30, 34*), so we explored multiple formats for the targeting ligands, including human 4-1BBL (h4-1BBL) in its native Type II transmembrane conformation (*35*), reformatted as a Type I transmembrane protein, and in multiple oligomerization states **(Fig. S1, Table S1)**. We additionally chose to display scFv-formatted versions of clinically evaluated anti-4-1BB agonists – urelumab and utomilumab – which display exquisite selectivity for human 4-1BB (*36*), and two additional agonists– TABBY 106 and TABBY 107 – which display cross-reactivity between murine and human 4-1BB proteins (*37*) **(Fig. S1, Table S1)**.

To compare transduction rates for each of these constructs, we used Jurkat T cell lines in which half of the cell population stably expresses either human or murine 4-1BB. Broadly tropic VSVGwt LV indiscriminately transduced 4-1BB+ and 4-1BB-Jurkat T cell populations, as expected given the ubiquitous expression of LDLR, the cognate receptor for VSVGwt (*38*). In comparison, each of the targeted lentiviral vectors co-displaying VSVGmut and a 4-1BB binder selectively transduced 4-1BB+ cells, albeit at different rates (**Fig. 1B-C, Fig. S1**). Viruses displaying high affinity urelumab or utomilumab scFvs transduced more potently than viruses displaying h4-1BBL, achieving >90% on-target transduction with minimal off-target or off-species transduction at the highest dose tested (**Fig. 1B-C, Fig. S1**). We calculated human or mouse 4-1BB-specific selectivity ratios for each candidate lentiviral construct and observed that lentiviruses displaying urelumab or utomilumab drastically increased transduction preference for human 4-1BB+ cells. Based on its efficient transduction and more favorable safety profile in previously conducted clinical studies as a single agent (*39, 40*) or in combination with Pembrolizumab (*41*), we selected the utomilumab-based construct as our lead 4-1BB-virus for subsequent experiments (**Fig. S1**). Collectively, these data demonstrate that lentiviruses can be specifically redirected to infect human 4-1BB-expressing T cells using reformatted agonistic scFvs.

### 4-1BB LVs specifically activate, expand, and transduce antigen-specific CD8 and CD4 T cells after peptide stimulation

We next determined if 4-1BB-virus could selectively transduce primary T cells that have recently encountered their cognate antigen. We established a model system using peptide restimulation of memory T cells specific for commonly encountered viral antigens, as these T cells are found at low abundances in most healthy donor peripheral blood mononuclear cells (PBMCs) (**Fig. 2A**). We initially focused on the peptide NLVPMVATV (NLV), which is an immunodominant epitope derived from cytomegalovirus (CMV) in HLA-A*02:01^+^ (HLA-A2^+^) individuals. Since 4-1BB expression reaches peak upregulation 24-48 hours after TCR stimulation and declines thereafter (**Fig. S2**), we stimulated PBMCs from HLA-A2^+^ healthy donors with NLV peptide, and added 4-1BB-targeted or untargeted (VSVGwt) lentiviral vectors encoding ZsGreen 24 hours later (**Fig. 2A**).

**Fig. 2.**
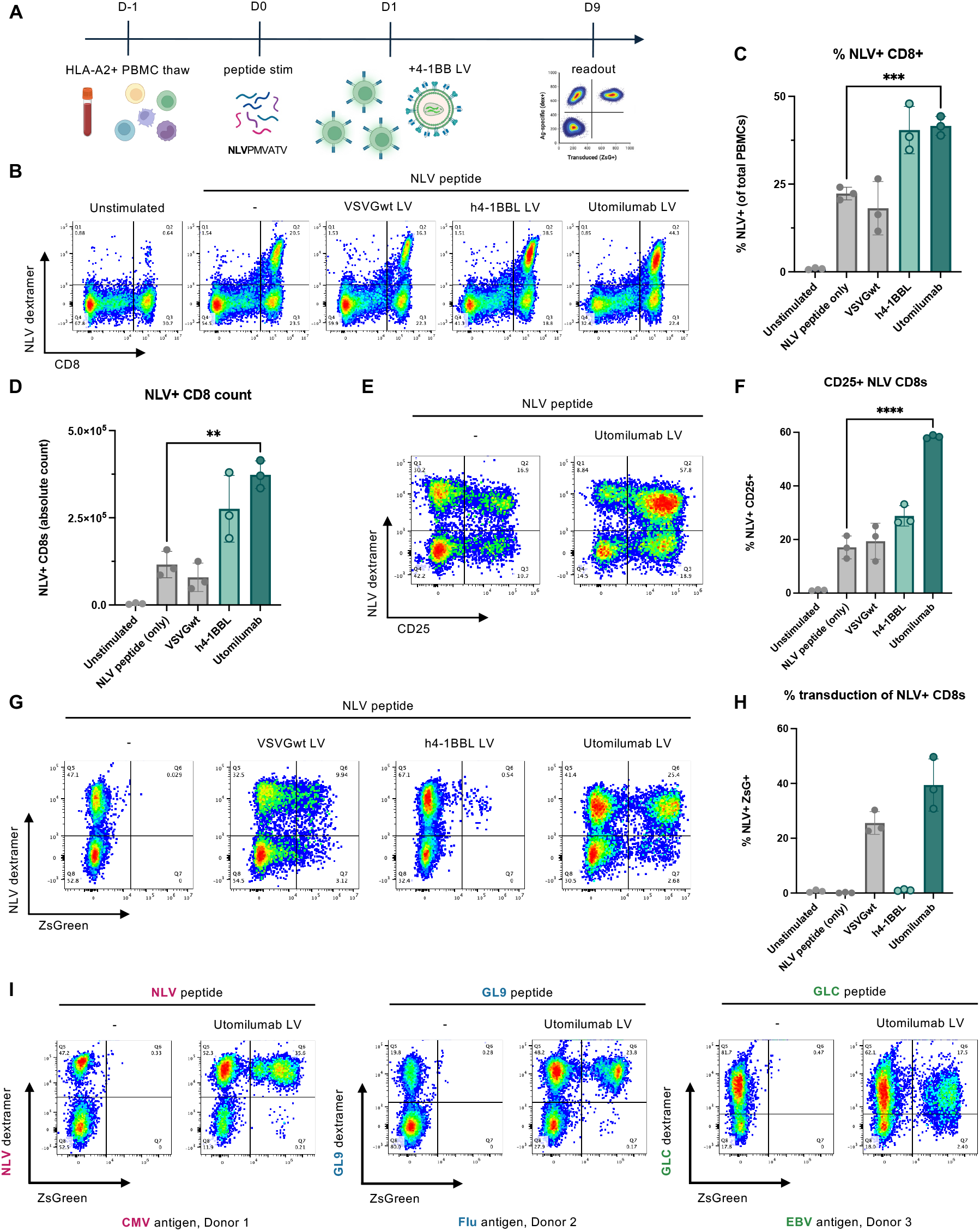
4-1BB LV specifically activates, expands, and transduces antigen-specific CD8 T cells after peptide stimulation, across multiple donors. **(A)** HLA-A2+ PBMCs are thawed and rested overnight (D-1) before peptides are added either in isolation or in pool (on D0), after which antigen-presenting cells within PBMC mixtures present MHC-restricted peptide(s) to cognate antigen-specific T cells. After 24 hours of peptide stimulation (D1), 4-1BB LV is added to PBMCs to target rare, recently-activated clonotypes within the pool of polyclonal cells via binding to 4-1BB on the surface of on-target, antigen-specific T cells, which can be readout via assays such as flow cytometry after several days of expansion (D9). Unless otherwise indicated, 4-1BB LVs package a ZsGreen fluorophore, enabling stable readout of previously-activated T cells. **(B)** Representative flow plots of 9-day expanded PBMCs gated on live cells (in the absence of dextran). “NLV dextramer” refers to multimerized HLA-A*02:01 presenting NLVPMVATV (CMV pp65) peptide to identify NLV-specific CD8+ T cells. **(C)** Quantitation of the percentage and **(D)** absolute count of NLV dextramer+ CD8+ T cells present after expansion. **(E)** Representative flow plots of PBMCs either stimulated with NLV peptide only or NLV peptide + Utomilumab LV, displaying activation (CD25+) in live+, CD8+ T cells. **(F)** Quantitation of the percentage of activated antigen-specific (CD25+ NLV+) CD8+ T cells after 9-day expansion in the presence of the indicated condition. **(G)** Representative flow plots of lead candidate LVs (h4-1BBL LV or Utomilumab LV) and control LV (VSVGwt), assessing transduction (ZsGreen+) in NLV dextramer+ T cells (gated on live+, CD8+). **(H)** Quantitation of transduction observed in antigen-specific NLV+ CD8s in (G). **(I)** Representative flow plots of PBMCs from 3 donors grown in the presence of peptide (NLV, GL9, or GLC) alone or with Utomilumab LV, assessing transduction (ZsGreen+) in NLV dextramer+ T cells (gated on live+, CD8+). Significance in (C), (D), and (F) was determined using a two-tailed unpaired Student’s t-test where \*\**P* <0.01, \*\*\**P* <0.001, and \*\*\*\**P* <0.0001.

Both transduced and untransduced antigen-specific cells showed greater expansion in the presence of the 4-1BB-targeted LVs, by frequency and absolute count, surpassing the expansion seen with NLV peptide only or the untargeted VSVGwt LV (**Fig. 2B-D**). Antigen-specific T cell expansion is likely mediated by the costimulatory effect of crosslinking 4-1BB on recently-activated T cells, as PBMCs exposed to NLV peptide and 4-1BB LV displayed greater CD25 expression in NLV-reactive T cells after nine days of expansion when compared to control cells exposed to NLV peptide only (**Fig. 2E-F**). Although both 4-1BBL and anti-4-1BB scFv viruses showed selective transduction of NLV dextramer+ T cells (**Fig. 2G**), utomilumab scFv virus achieved approximately 40-fold higher transduction of target cells than viruses displaying h4-1BBL (**Fig. 2H**).

To establish that our approach could generalize to other antigens, we stimulated three different healthy donor PBMC samples with peptides from common viruses – GL9 (derived from influenza A) and GLC (derived from Epstein-Barr virus) in addition to NLV – followed by 4-1BB LV and observed expansion and selective transduction in distinct antigen-specific CD8+ T cells (**Fig. 2I**). We repeated the PBMC stimulation protocol using a pool of 95 class I MHC-restricted peptides derived from commonly-encountered antigens (‘CEF’), as well as 35 peptides presented by class II MHCs for recognition by CD4+ T cells (‘CEFTA’), to observe whether the approach is capable of transducing multiple antigen reactivities simultaneously. We observed selective transduction and expansion of CD8+ and CD4+ T cells in the presence of both CEF and CEFTA peptide pools and 4-1BB-virus, respectively, as compared to cells stimulated with peptide pools without 4-1BB lentivirus addition (**Fig. S3-4**). Importantly, we observed NLV-reactive T cell transduction only when NLV peptide was added to the large class I MHC peptide pool (‘95 CEF’) or class II MHC peptide pool (‘CEFTA + NLV’), but not when the NLV peptide was absent from the pool (‘94 CEF-NLV’ or ‘CEFTA’ alone), indicating that 4-1BB virus transduces cells after cognate antigen-driven TCR activation, and can do so amongst a pool of other expanding CD8+ or CD4+ reactivities (**Fig. S3-4**). CD8+ T cells displayed sustained activation and proliferation in the presence of NLV peptide stimulation and 4-1BB virus, both when NLV was added as a single peptide or in a pool of peptides (**Fig. S3**). Taken together, these data highlight the ability of 4-1BB LV to specifically target rare, recently-activated 4-1BB+ CD4 and CD8 T cells.

### 4-1BB virus faithfully transduces antigen-specific CD8+ T cells, preserving TCR clonotypic diversity during expansion

While 4-1BB virus expands antigen-specific T cells, only a subset of the expanded cells are transduced. Transduction of antigen-specific clones could be stochastic, or TCR affinity of clonotypes could influence their transduction rate. To distinguish between these two possibilities, we used single cell TCR sequencing to define the clonal repertoire of transduced versus non-transduced tetramer-positive T cells.

PBMCs from two HLA-A*02:01+ healthy donors were stimulated with two pooled viral peptides (NLV and GL9) (**Fig. 3A**). Addition of 4-1BB virus led to a significant increase in expansion of NLV and GL9 tetramer-positive cells in each donor; in Donor 1 both NLV and GL9 reactive T cells expanded, whereas expansion in Donor 2 was largely comprised of NLV-reactive CD8+ T cells **(Fig. S5)**. After two weeks of expansion, cells were labeled with barcoded antibodies to CD4 and CD8 as well as barcoded hashtag antibodies and tetramers prior to sorting as per the gating scheme in **Fig. S5**. TCR sequencing revealed a heavily skewed distribution of TCR clonotypes **(Fig. 3B)**, indicative of a large clonal expansion in both donors across all of the experimental groups combined. In Donor 1, where we observed significant NLV tetramer+ and GL9 tetramer+ CD8+ T cell expansion, we identified an oligoclonal set of uniquely expanded TCR clonotypes, whereas in Donor 2 where we observed greater expansion of NLV tetramer+ CD8+ T cells we detected a nearly monoclonal response, with only 2 clonotypes representing nearly all sequenced TCRs in this donor **(Fig. 3B, Fig. S6)**.

**Fig. 3.**
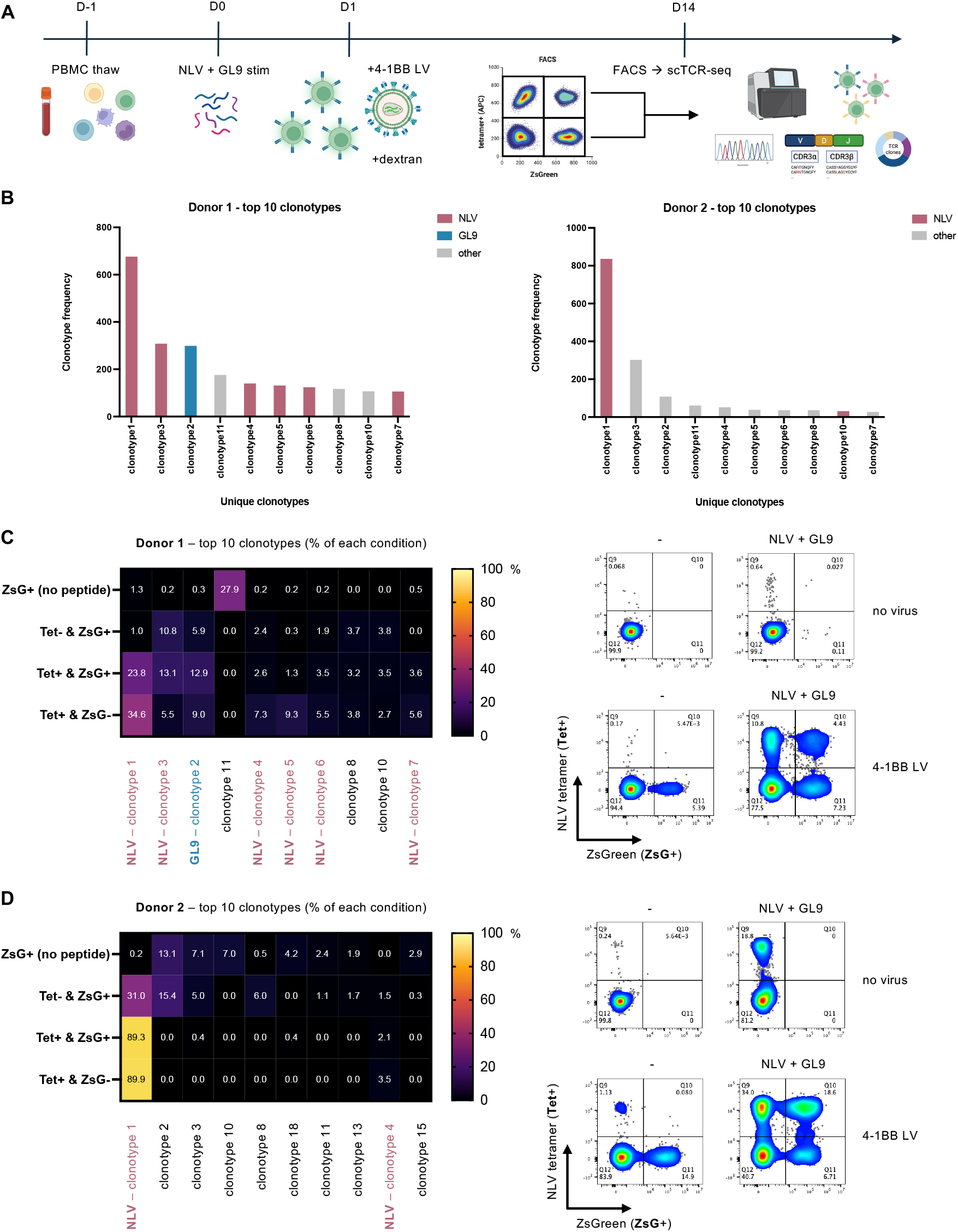
4-1BB LV faithfully transduces antigen-specific CD8+ T cells, preserving TCR clonotypic diversity during expansion. **(A)** Schematic of experiment workflow in which HLA-A2+ PBMCs were thawed on D-1, rested overnight, stimulated with NLVPMVATV (NLV) + GILGFVFTL (GL9) peptides on D0, after which 4-1BB LV was added 24 hours later in the presence of dextran transduction enhancer. After two weeks of expansion (D14), T cells were individually sorted via FACS and sequenced with scTCR-seq. **(B)** Histograms of the top 10 uniquely expanded clonotypes in donor 1 (left) and donor 2 (right). Clonotypes in donor 1 and donor 2 are highlighted by antigen specificity (NLV, GL9, or “other” indicating neither NLV nor GL9 barcode detection). **(C)** Left: Heatmap of the top 10 expanded clonotypes in donor 1. Numbers indicate the relative proportion of uniquely sequenced clonotypes corresponding to the indicated clonotype (column) per condition (row). Clonotypes are listed with arbitrary numbers, however each clonotype is labeled with the detected antigen specificity (NLV, GL9, or unlabeled for “other”). Right: day 14 PBMC expansions prior to sorting for single cell TCR sequencing. Representative flow plots of PBMCs grown in the absence (-) or presence of indicated peptides (NLV + GL9 peptide pool), in the absence (no virus) or presence of 4-1BB LV with dextran. Conditions were assessed for NLV-specific expansion (NLV tetramer^+^) and transduction (ZsGreen^+^) prior to scTCR-seq, gated on live cells. **(D)** Same analysis as in (C) but for donor 2.

When examining the top 10 expanded clonotypes across all groups, subdivided by sorted experimental condition, we observed that the top 5 expanded clonotypes in the tetramer-positive sorted conditions in Donor 1 did, indeed, correspond to the input antigen specificities (NLV & GL9) based on sequencing of barcoded tetramers **(Fig. 3C, Fig. S6, see Materials and Methods)**. This observation held for Donor 2, as well, with ∼90% of all observed T cells in the tetramer-positive conditions corresponding to a single NLV-reactive clonotype (“clonotype 1”) **(Fig. 3D)**. When compared to conditions in which no peptide was added, the ZsG^+^ (transduced) TCR clonotypes were mutually exclusive with those that were present in the tetramer-positive sorted conditions, indicating that these transduced clonotypes were not reactive to either NLV nor GL9 **(Fig. 3C-D, Fig. S6)**. We compared the top 10 expanded CDR3beta and CDR3alpha sequences per donor to VDJdb (*42*) – a curated database of published TCRs with known antigen specificities – in an attempt to both validate identified clonotypes and to uncover newfound reactivities. Interestingly, we detected two matches in Donor 1 for CDR3s contained within clonotypes 2 and 4 and the corresponding antigen specificities of GL9 and NLV, respectively, which directly matched what we detected by scTCR-seq of the barcoded GL9 and NLV tetramers we included in the experiment, corroborating our results and supporting the finding that 4-1BB LV is capable of transducing *bona fide* antigen-specific public clonotypes found in the population (**Fig. S6**).

Importantly, we observed substantial overlap with the same dominant clonotypes sequenced in the tetramer^+^ ZsG^+^ (transduced) and tetramer^+^ ZsG^-^ (untransduced) groups (**Fig. 3C-D**). These data fit with the model that 4-1BB LV provides a costimulatory signal to 4-1BB+ T cells at the time of coculture (24 hours after antigen stimulation) to promote expansion (as demonstrated in Fig. 2), while transducing a subset of the same responding clonotypes. Together, these data demonstrate that we are able to successfully sort, sequence, and identify expanding T cell clones while simultaneously uncovering T cell lineage and antigen specificity on a per-cell basis from multiple donors.

### BB LV can successfully deliver therapeutic cargoes to antigen-specific cells *in vitro* and control antigen positive human melanoma *in vivo*

With the ability to successfully expand and transduce antigen-specific T cells from rare starting populations, we next introduced immunomodulatory genetic cargoes into these cells via 4-1BB LV. To create a model to functionally test these cargoes, we engineered tumor cells to express a model antigen (NLV) linked to the C-terminus of the fluorophore mScarlet for proof-of-concept studies recognized by readily-available, viral antigen-reactive T cells (**Fig. S7**). For our initial experiments, we chose to transduce the A375 cell line with the mScarlet-NLV construct, as these cells endogenously express HLA-A*02:01 at baseline, and A375 human melanoma represents a challenging solid tumor model that could more accurately mimic intratumoral dynamics, compared to *in vitro* experiments (*43-45*). Given the hostile tumor microenvironment in solid tumor settings, we identified genetic cargoes which could offer pro-survival signals to enhance intratumoral T cells. We chose a membrane-tethered version of the computationally-designed IL-2/IL-15 mimetic (called Neo2/15), as this cytokine is profoundly stable and possesses high affinity for IL-2Rβ γ_c_ (*46*), providing a potent signal to antigen-specific T cells. We also chose to express a Fas switch receptor – Fas-4-1BB – as it offers the possibility of turning a pro-apoptotic signal (FasL-trigged cell death) into a pro-survival signal (4-1BB signaling), especially within the TME (*47*) (**Fig. S7**).

We transduced healthy donor PBMCs 24 hours after NLV peptide stimulation with either Neo2-ZsGreen or Fas-4-1BB-ZsGreen cargoes and assessed transduction (via ZsGreen expression) and anti-tumor activity *in vitro* after two weeks of expansion. When the engineered T cells were co-cultured with A375 cells expressing NLV (mScarlet^NLV^), we observed robust *in vitro* cell killing, whereas A375 cells engineered to express ELAGIGILTV, an HLA-A2-restricted control peptide, (mScarlet^ELA^) were not affected (**Fig. S7**). NLV peptide-expanded but untransduced CD8+ T cells were able to similarly lyse on-target A375-mScarlet^NLV^ tumor cells, but not off-target A375-mScarlet^ELA^ tumor cells, however, they did so with slower kinetics than either the Neo2-ZsGreen or Fas-4-1BB-ZsGreen cargo-containing CD8+ T cells (**Fig. S7**).

We next expanded healthy donor PBMCs with NLV peptide in the presence or absence of 4-1BB LV particles packaging either Neo2-ZsGreen or Fas-4-1BB-ZsGreen cargoes prior to adoptive transfer into A375-mScarlet^NLV^ tumor-bearing NSG mice (**Fig. 4A and S8**). After two weeks of expansion, we observed a significant increase in the percentage of NLV+ CD8+ T cells in PBMCs exposed to cargo-containing 4-1BB LVs, compared to NLV peptide alone, and relative to unstimulated PBMCs. Within the PBMCs receiving 4-1BB LV, we observed significant transduction within the antigen-specific T cell compartment: 38% (Neo2) and 55% (Fas-4-1BB) of total NLV+ CD8+ cells (**Fig. 4B**). Seven days post-inoculation of A375-mScarlet^NLV^ tumor cells, mice either received PBS (mock), 5×10^6^ NLV-reactive T cells expanded with NLV peptide only, or with 4-1BB LV packaging either Neo2 or Fas-4-1BB genetic cargoes. We observed several complete responses in mice receiving NLV+ CD8+ T cells engineered with Neo2-ZsGreen or Fas-4-1BB-ZsGreen, but not those adoptively transferred with NLV+ CD8+ T cells expanded with NLV peptide alone (**Fig. 4C-D**). These responses were durable, particularly in the Neo2-ZsGreen cohort, in which half of mice remained tumor-free at study endpoint (day 52 post-ACT). Collectively, these data indicate that the tumor growth delay and eradication observed in mice receiving Neo2-ZsGreen or Fas-4-1BB resulted in significant survival extension compared to mock and NLV peptide only groups (**Fig. 4E**).

**Fig. 4.**
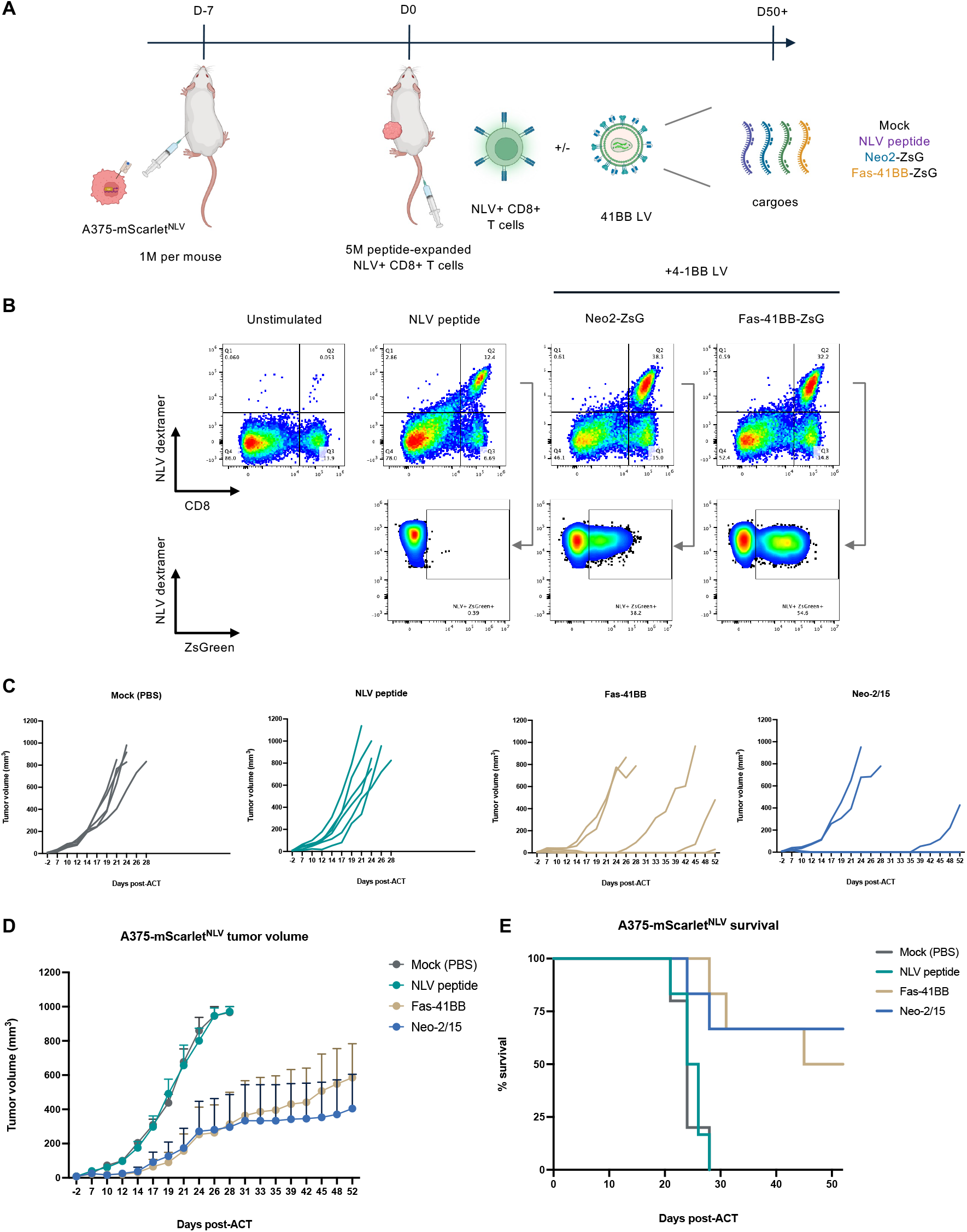
4-1BB LV delivery of immunomodulatory cargoes significantly delays tumor growth and extends survival in a xenograft melanoma model. **(A)** Schematic of the study design in which NSG mice initially received 1×10^6^ A375-mScarlet^NLV^ tumor cells in the flank, after which antigen-specific T cells were adoptively transferred via tail vein injection. After 7 days of engraftment, 5×10^6^ peptide-expanded NLV+ CD8+ effector T cells were transferred per mouse and the experimental cohorts in this study were: 1) Mock (PBS-treated), 2) NLV peptide only, 3) NLV peptide + 4-1BB LV containing Neo2-ZsGreen, and 5) NLV peptide + 4-1BB LV containing Fas-41BB-ZsGreen. **(B)** Representative flow plots of infusion product PBMCs that were unstimulated or stimulated with NLV peptide +/- the indicated cargo groups. Top: assessing expansion of antigen-specific NLV+ CD8+ effector T cells. Bottom: Assessing transduction within the antigen-specific NLV+ CD8+ T cell compartment. **(C)** Individual tumor curves plotting tumor volume (mm^3^) over time, represented as days after adoptive cell transfer (ACT). **(D)** Tumor volume curves represented as mean +/- SEM (N=5 for mock (PBS) treated mice, N=6 mice per experimental condition). **(E)** Kaplan-Meier curve representing overall survival for the length of the experiment after ACT.

### 4-1BB LV enables transduction of patient-derived TILs

Our results through this study represented antigen-specific transduction within healthy donor samples and thus we next sought to apply targeted lentiviral vectors to tumor-derived samples from patients with metastatic cancer, as these primary samples carry additional challenges of T cell viability, refractory activation, skewed cell phenotypes, and difficulty with expansion and transduction (*48-50*).

To do so, we began by exposing cryopreserved tumor-associated lymphocytes derived from the ascites fluid of a patient with ovarian cancer to 4-1BB LV 24 hours after anti-CD3/anti-CD28 stimulation. These patient samples include autologous tumor cells, and thus antigens presented on MHC should be able to induce activation of antigen-specific T cells. After 7 days of expansion, ∼5% of T cells expressed ZsGreen, indicating that T cells from immunosuppressive ascites are capable of transduction via 4-1BB **(Fig. 5A)**. We next turned to using fresh TIL from tumor fragments isolated from resected ovarian cancer. After two days in culture, individual TIL fragments were independently assessed for 4-1BB expression. As shown in **(Fig. 5B)**, the percentage of CD3+ 4-1BB+ T cells ranged from 10-30% across all 24 plated tumor fragments, likely representing spatial heterogeneity of T cell clones present within tumor tissue (*51*). We selected 15 of these tumor fragments for further study based on cell count, viability, and T cell 4-1BB expression and tested increasing doses of 4-1BB LV expressing ZsGreen to assess transduction in otherwise unstimulated TIL **(Fig. 5C)**. As before, we observed a dose-dependent increase in ZsGreen signal from ∼5% at the lowest dose tested (1.25 µL) to ∼25% at the highest dose (10 µL) (**Fig. 5D, Fig. S9**). When plotted against initial 4-1BB expression (on day 2) per matched tumor fragment, we observed a dose-dependent increase in the level of transduction per ‘transducible’ (4-1BB+) cells **(Fig. 5E)**, indicating reproducible on-target transduction of fresh TIL, across different tumor fragments.

**Fig. 5.**
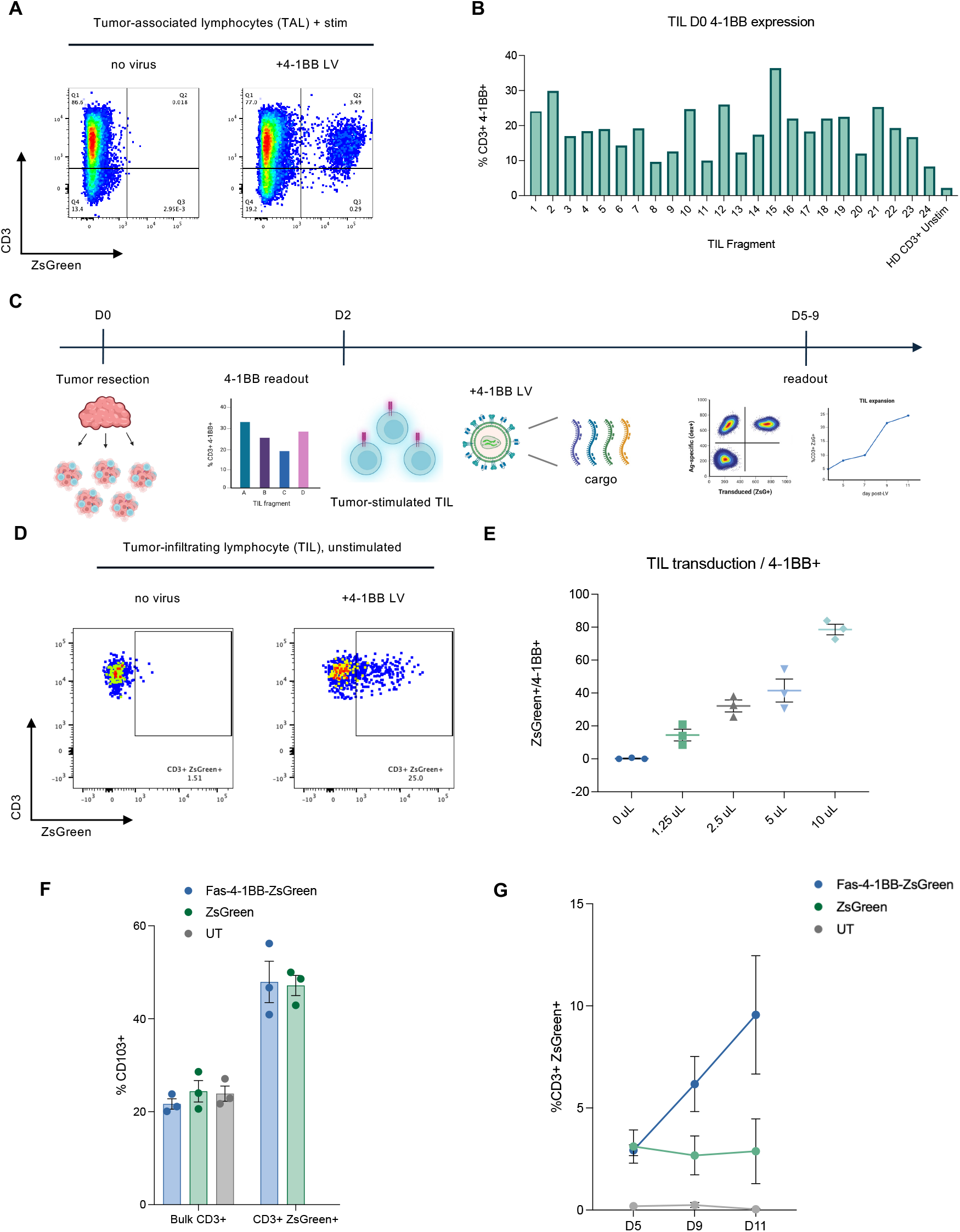
4-1BB LV is capable of specifically transducing primary TIL samples *ex vivo*. **(A)** TAL samples that were pre-activated with anti-CD3/anti-CD28 prior to 4-1BB LV addition were assessed for transduction in CD3+ T cells after 7 days. **(B)** Bar plots summarizing the percentage of CD3+ 41BB+ T cells from 24 different tumor fragments derived from a single surgical resection of an ovarian tumor with healthy donor (“HD”) CD3+ unstimulated T cells included as a negative control. **(C)** Schematic of TIL workflow using a fresh tumor resection from a patient with ovarian cancer on D0, after which time TIL within individual tumor fragments were grown in the presence of IL-2 until D2 when 4-1BB+ expression is measured within the CD3+ T cell compartment. On D2, 4-1BB LV containing either ZsGreen only or Fas-4-1BB-ZsGreen cargo are added to tumor-stimulated TIL and allowed to expand until D5-D9 when TIL are assessed via flow cytometry for transduction, phenotype, and count. **(D)** Representative flow cytometry plots of TIL unexposed to LV (“no virus”) or 10 µL of 4-1BB LV containing ZsGreen. **(E)** Triplicate tumor fragments were exposed to four different doses (10, 5, 2.5, 1.25 µL) of 4-1BB LV or no virus (0 µL) and transduced TILs (ZsG^+^) at day 2 were compared to their matched, baseline 4-1BB+ expression at day 0; the fraction of ‘transducible’ TIL are represented as the percentage of ZsGreen+/41BB+ TIL. **(F)** Bar plots representing the percentage of CD103+ (tumor-reactive) TIL within the CD3+ compartment from either bulk CD3+ T cells or CD3+ T cells that were transduced (ZsGreen+) with either 4-1BB LV containing ZsGreen only or Fas-4-1BB-ZsGreen cargo virus on day 5; “UT” represents untransduced TIL not exposed to 4-1BB LV. **(G)** Line plots representing the percentage of transduced TIL (CD3+ ZsGreen+) over time (days 5, 9, and 11) after exposure to 4-1BB LV containing ZsGreen or Fas-4-1BB-ZsGreen or no LV, untransduced (UT).

Encouraged by these data, we wondered if 4-1BB LV could be used to deliver immunomodulatory cargo to fresh TIL. We obtained a metastatic lesion resected from the omentum of a separate patient with ovarian cancer, isolated tumor fragments from within this resected mass, and plated them in culture with high dose IL-2. As before, we assessed the 4-1BB expression of each tumor fragment after two days in culture and observed 3-15% expression across all 40 fragments tested (**Fig. S9**). We then moved forward with a subset of these fragments and left them untransduced or exposed them to either 4-1BB LV containing ZsGreen or Fas-4-1BB-ZsGreen virus. After 5 days in culture with virus, we observed ∼8% transduction of CD3+ cells exposed to ZsGreen-containing 4-1BB LV and ∼3% transduction of CD3+ cells exposed to Fas-4-1BB-ZsGreen-containing 4-1BB LV at the top doses tested, representing ∼70% and ∼40% of matched 4-1BB+ CD3+ cells on day 2, respectively (**Fig. S9**). Despite representing only a subset of all cells, the transduced CD3+ TIL expressed significantly more CD103+ when compared to untransduced cells, which correlates with increased tumor-reactivity within the TIL compartment (*52*) (**Fig. 5F**). Lastly, when these transduced TIL were allowed to expand in culture over time, the CD3+ TIL transduced with Fas-4-1BB ZsGreen preferentially proliferated relative to CD3+ TIL transduced with ZsGreen only (**Fig. 5G**), suggesting that 4-1BB LV delivery of genetic cargo can drive a distinct pro-survival phenotype in fresh TIL derived from patients with metastatic cancer.

## DISCUSSION

Here, we demonstrate a feasible, scalable approach to target recently-activated T cells via 4-1BB using engineered LV particles. This platform provides direct costimulation to antigen-specific CD4 and CD8 T cells via 4-1BB crosslinking, while simultaneously introducing immunomodulatory cargoes that can alter cell fate. In resting cells, 4-1BB is expressed at low levels, but its expression after T cell activation is temporally controlled in a manner that offers superb signal-to-noise (*28*), and can capture multiple distinct reactivities in a pooled format (*53*). In doing so, our LV approach enables simultaneous expansion, transduction, and engineering of rare antigen-specific T cells from primary cell peripheral blood or tumor-derived polyclonal T cells without the need for *a priori* knowledge of peptide antigen or the responding MHC.

Our approach to target recently-activated T cells via 4-1BB using engineered LV particles can enrich antigen-specific T cells from polyclonal cell populations, across a range of starting frequencies. In doing so, we envision this platform being applied to both patient-specific TIL and peripheral blood samples given the commonality of 4-1BB expression after TCR cross-linking, regardless of antigen source (*54-56*). One could envision *in vivo* applications of 4-1BB LV as an intratumoral adjuvant or systemic delivery (*57*) after an antigen-priming event such as vaccination or clonotype reinvigoration with immune checkpoint blockade. These approaches would enable therapeutic delivery of immunomodulatory cargo transfer directly to endogenously-generated antigen-specific T cells. For research purposes, we can use 4-1BB LV to transduce responding clonotypes and sequence their TCRs using routine scTCR-seq pipelines. The ability to directly track responding clonotypes *in vitro* and *in vivo* offers the unique possibility of understanding both the diversity and temporal dynamics of T cell antigen specificity from various HLA haplotypes and sample sources, all using a single reagent.

The modular nature of LV surface engineering enables facile development of receptor-targeted viral vectors with the ability to infect specific cell types (*30*). However, the exact choice of the best individual ligand or scFv architectural format still requires empiric testing per intended target. For instance, utomilumab does not possess the highest affinity for 4-1BB receptor relative to the other scFvs tested, and type I TM formatted ligands outperformed the native type II TM formatted ligands, indicating that native ligand architecture is not necessarily the most ideal format to engage with a target receptor when presented on the surface of a LV particle. Whether the efficacy of the utomilumab-presenting 4-1BB LV is driven by the robust scFv expression observed on producer cells, its nanomolar affinity for 4-1BB, or the fact that multimerized scFvs on the LV particle surface might offer an avidity advantage for receptors like the TNF-family receptor protein 4-1BB which requires trimerization (*58*) for successful activation and internalization remains to be examined. Future experiments are needed to further elucidate ‘rules’ of this LV targeting platform and whether specific receptors or receptor families have a greater propensity for endocytosis-driven transduction.

In this work we applied the 4-1BB LV platform to anti-viral T cells in an *in vivo* model of human melanoma and in an *ex vivo* demonstration of TILs using autologous tumor as the antigen source for 4-1BB upregulation in a manner agnostic to both peptide and MHC. We envision using this approach for enhance adoptive T cell engineering and TCR discovery efforts, but also with wide applicability beyond cancer applications, as 4-1BB expression is a common pathway shared by T cells after cognate antigen exposure (*59*). Given the easily portable and scalable nature of 4-1BB LV, one could imagine use cases in selectively identifying and tracking T cell responses in infectious disease applications or modulating T cells in autoimmune diseases. Moreover, this work demonstrates the modularity of surface-engineered LV vectors, allowing for facile development of other receptor-targeting LVs with cell-specific infection capability. Future engineered LVs could even be used in combination alongside 4-1BB LV, targeting multiple arms of innate and adaptive immunity, serving as adjuvants to better coordinate ongoing immune responses. Further efforts can use these engineered 4-1BB LVs as off-the-shelf vectors to achieve therapeutic effect, uncover novel TCRs, and even expand rare antigen-specific T cells all with the use of a single, engineered viral particle.

## MATERIALS AND METHODS

### Plasmids

psPAX2 (plasmid #12260), pHIV-ZsGreen (plasmid #18121), and pMD2.G (VSVGwt, plasmid #12259) were obtained from Addgene. pMD2-VSVGmut was developed in the Birnbaum lab (*30*) and is publicly available on Addgene (plasmid #182229). Human 4-1BBL (Uniprot P41273) and murine 4-1BBL (Uniprot P41274) constructs were designed by us and obtained from Twist Biosciences and cloned into the pMD2 backbone. Utomilumab scFv sequence was obtained from Patent US 2013/0078240 A1, urelumab scFv sequence was obtained from Patent US 2012/8137667 B2, and TABBY 106 and TABBY 107 scFv sequences were obtained from Patent WO 2017/205745 A1. Gene blocks for all 4-1BB-targeting constructs displayed in **Fig. S1** and listed in **Table S1** were ordered from IDT and cloned into the pMD2 backbone. pBS_ZsGreen was synthesized by cloning ZsGreen from pHIV-ZsGreen and transferring into a pLenti backbone (plasmid #181970 from Addgene). Full-length human 4-1BB (Uniprot Q07011) and murine 4-1BB (Uniprot P20334) were ordered as gene blocks and cloned into a pHIV-TCR-IRES-tEGFR backbone from the Birnbaum lab by first excising the TCR prior to Gibson Assembly (New England Biolabs #E2611). pHIV-mScarlet^NLV^ was designed by taking the full-length mScarlet fluorophore sequence from Addgene plasmid #85042 and placing a flexible C-terminal GSG linker followed by a 17aa sequence for NLV (pp65) or an 18aa sequence for ELA (MART-1), both of which contain the immunogenic 9-mer (NLVPMVATV) and 10-mer (ELAGIGILTV) peptides, respectively, flanked by 4aa of native peptide sequence at the N- and C-termini of each (see **Fig. S7** for more). Both constructs were ordered as a gene block from IDT and cloned via Gibson Assembly into the aforementioned pHIV-IRES-tEGFR backbone. pBS_Neo2-P2A-ZsGreen and pBS_Fas-4-1BB-P2A-ZsGreen cargo plasmids were ordered as gene blocks and cloned into the pBS_ZsGreen cargo backbone (described in **Fig. S7**). Full annotated sequences for all constructs can be found in **Tables S1-2**.

### Cell lines

Jurkat (clone E6-1), A375, and HEK-293-T cells were all purchased from ATCC. Jurkat cells were cultured in RPMI-1640 (ATCC) + 10% FBS + 1% pen-strep. A375 and HEK-293-T cells were cultured in DMEM (ATCC) + 10% FBS + 1% pen-strep. All cells were maintained in routine tissue culture incubators (37C, 5% CO_2_). Jurkat T cells for initial 4-1BB LV screening were transduced with pHIV-h4-1BB-IRES-tEGFR or pHIV-m4-1BB-IRES-tEGFR (described in “Plasmids” above) and sorted on h4-1BB or m4-1BB expression, respectively, at ∼50:50% 4-1BB+:4-1BB-expression to intentionally preserve 4-1BB-Jurkat T cells as internal “off-target” control cells. In experiments using human or murine 4-1BB-expressing Jurkat T cells, EGFR is used as a surrogate marker for 4-1BB expression (unless otherwise stated), as EGFR expression and detection is orthogonal to the entry of 4-1BB-targeting LV vectors. A375 were transduced with pHIV-mScarlet-GSG-NLV-IRES-tEGFR or pHIV-mScarlet-GSG-ELA-IRES-tEGFR and sorted on mScarlet^hi^ cells three times to achieve purity in order to generate the A375-mScarlet^NLV^ and A375-mScarlet^ELA^ stable cells lines, respectively.

For the experiment in **Fig. 1**, h4-1BB Jurkat T cells were plated in flat-bottom 96-well plates at a density of 0.5×10^6^ cells/mL in 100 µL of RPMI-1640 + 10% FBS + 1% pen-strep complete media with dextran (final concentration = 0.8 µg/mL), unless otherwise stated. Candidate 4-1BB LVs were then added in a dilution series of 1, 0.25, 0.0625, and 0.015625 µL (in triplicate) in 100 µL of complete RPMI-1640 media. Cells + LV mixtures were gently pipetted up and down and placed in a humified incubator at 37C, 5% CO_2_ and left undisturbed for 48 hr prior to assessing for transduction efficiency via flow cytometry. LV titers were calculated using the following formula:

LV titer (TU/mL) = [(plated target cell #)*(ZsGreen% transduction)]/(viral volume in µL/1,000)

### Peripheral blood mononuclear cells (PBMCs)

All PBMCs used in this work were isolated from leukapheresis collections obtained from StemCell Technologies or StemExpress (now CGT Global) and processed using the EasySep Direct Human PBMC Isolation Kit (StemCell Technologies #19654). Following isolation, all PBMCs were cryopreserved at 50-100 x 10^6^ cells/vial in 90% FBS + 10% DMSO and stored in liquid nitrogen until future use for experimentation.

For peptide expansions described in **Figs. 2-4**, PBMCs were thawed and rested overnight in RPMI-1640 (ATCC) + 10% FBS + 1% pen-strep at a density of 1-2 x 10^6^ cells/mL in a humified incubator at 37C, 5% CO_2_. On the following day, PBMCs were collected and live cells were plated with or without peptide at 1.5 x 10^6^ cells/mL in 200 µL per well of a 96-well U-bottom plate. All peptides were used at 1 µg/mL final concentration unless otherwise specified. All MHC class I peptides used in this study were purchased from GenScript at crude purity. MHC class II ‘CEFTA’ peptides were purchased from Mabtech (PepPool #3617-1) and were prepared and used at a final concentration of 2 µg/mL per manufacturer’s instructions. All peptides used in this study are listed in **Table S3**.

Twenty-four hours after peptide stimulation, 100 µL of media was carefully removed off the top of 96 U-bottom wells without disturbing the clustered PBMCs and decanted. This media was replaced with 100 µL of virus-containing media (+/- dextran, depending on the experiment) and the virus + cell mixture was pipetted gently up and down before being placed back in the incubator for expansion. If dextran was used, it was added to virus-containing media at a final concentration of 0.8 µg/mL. 4-1BB LV was added to PBMCs at a multiplicity of infection (MOI) = 2, unless otherwise stated. Titer (TU/mL) calculations were made per batch of concentrated virus using h4-1BB Jurkats (as described above) to standardize across LV vectors. Antigen-specific T cells were grown in 96-well U-bottom plates until reaching confluence (typically ∼day 9-12, depending on the donor) throughout which time cells were supplemented with RPMI-1640 + 10% FBS + 1% pen-strep + 50 IU of human IL-2 (R&D systems). After reaching confluence, cells were either split 1:2 for flow cytometry evaluation on day 14, or pooled per condition, recounted, and replated at a density of 1 x 10^6^ cells/mL if expanding for indicated experiments. Calculation of MOI:

MOI = [viral titer (TU/mL)*virus volume (mL)]/(PBMC cell #)

For the experiments in **Figs. S3-S4**, PBMCs were initially labeled with CellTrace Violet Cell Proliferation Kit (ThermoFisher Scientific #C34557) at 5 µM per the manufacturer’s instructions, plated at 1.5 x 10^6^ cells/mL in 200 µL per well of a 96-well U-bottom plate and stimulated with the indicated peptides, as above. Utomilumab antibody was purchased from ThermoFisher Scientific (cat #MA5-42125) and added to PBMCs at a final saturating concentration of 1 µg/mL (*36*), unless otherwise stated.

For the experiment in **Fig. S2**, PBMCs were plated at 1 x 10^6^ cells/mL in complete RPMI-1640 without IL-2 and either left unstimulated or activated with the indicated volume of Gibco Dynabeads Human T-Activator CD3/CD28 (ThermoFisher Scientific #1131D) – 10 or 25 µL – per 1 million PBMCs. Triplicate conditions were assessed for 4-1BB+ expression in CD4+ and CD8+ T cells via flow cytometry at the indicated timepoints. After 4 days of stimulation (“d4”, indicated by the arrow), half of the experimental conditions had anti-CD3/CD28 Dynabeads magnetically removed and were allowed to continue growth in the incubator while continuing to be monitored for 4-1BB expression. These conditions – termed ‘d4 rest’ – were compared to the remaining half of experimental cells left either unactivated or continually activated with either 10 or 25 µL of sustained Dynabead (DB) stimulation.

### Animals

Female NOD/SCID/IL2R^null^ (NSG) mice were housed in the Koch Institute facility under the guidelines of MIT’s Division of Comparative Medicine and the MIT Committee on Animal Care. NSG mice were bred in-house and used in experiments described in **Fig. 4** and **Fig. S8** at 8-12 weeks of age.

T cell dosing pilot study sought to assess the number of NLV+ CD8+ T effector cells required to demonstrate efficacy in a flank tumor model harboring 1×10^6^ A375-mScarlet^NLV^ tumor cells (**Fig. S8**). We observed a dose-dependent decrease in tumor volume with increasing doses of NLV+ CD8+ T cells adoptively transferred after 7 days of tumor engraftment, and we chose to pursue the 5×10^6^ dose for further study in **Fig. 4**, in which there was a small efficacy observed with NLV peptide-expanded cells to mimic the efficacy of current TIL approaches.

For the experiment represented in **Fig. 4**, 1×10^6^ A375-mScarlet^NLV^ tumor cells were injected subcutaneously into the flank of NSG mice. After 7 days of engraftment, mice were randomized into various experimental groups prior to receiving 5×10^6^ NLV+ CD8+ T cells via tail vein for adoptive cell therapy (ACT). Tumors were measured 2-3 times per week by caliper measurements and tumor volume was assessed using the following formula: V = ½ (L*W^2^), with W representing the shortest dimension. Mice were monitored for signs of distress, weight loss, and tumor ulceration and euthanized at humane endpoint or when tumor volume surpassed 1000mm^3^.

### Lentivirus production

HEK-293-T cells were grown to 80-90% confluence in T-225 flasks, at which point they were transfected with the following DNA plasmids using TransIT-Lenti transfection reagent (Mirus Bio #MIR 6600):

42 µg transfer/cargo plasmid

22.5 µg psPAX2 helper plasmid

7.5 µg VSVGmut (or VSVGwt)

22.5 µg targeting plasmid (anti-4-1BB)

Viral supernatant was collected after 48 and 72 hr, pre-cleared for 10 min at 4C at 1,000 *xg* to remove sticky cell debris, prior to ultracentrifugation using a Thermo Scientific Sorvall LYNX 6000 for 90 min at 4C at 100,000 *xg*, after which viral supernatant was gently discarded and viral pellets were solubilized overnight at 4C in Opti-MEM prior to being aliquoted and frozen down at -80C for long-term use. Concentrated LV batches typically achieved 200-250x concentration from initial unconcentrated volume.

#### Jurkat screen calculation

For the bar plots displayed in **Fig. 1** and **Fig. S1**, calculations were obtained as follows:

#### h4-1BB+/h4-1BB-transduction ratio calculation

(4-1BB+ ZsGreen+ / ((4-1BB+ ZsGreen+) + (4-1BB+ ZsGreen-))*100 = On-target transduction % (4-1BB-ZsGreen+ / ((4-1BB-ZsGreen+) + (4-1BB-ZsGreen-))*100 = Off-target transduction % h4-1BB+/h4-1BB-transduction ratio = On-target transduction % / Off-target transduction %

#### h4-1BB/m4-1BB selectivity ratio calculation

human 4-1BB+ ZsG+ % / murine 4-1BB+ ZsG+ % per matched virus dose.

#### Barcoded tetramer production

Recombinant HLA-A2 heavy chain fused to a C-terminal AviTag and β_2_-microglobulin were expressed in *E. coli*, isolated as inclusion bodies, and solubilized in 8 M urea. The heavy chain was refolded in the presence of a three-fold molar excess of β_2_-microglobulin and a ten-fold molar excess of synthetic peptide (NLVPMVATV or GILGFVFTL) in 100 mM Tris-HCl (pH 8.0) supplemented with 400 mM L-arginine, 2 mM EDTA, 3 mM cysteamine, 3.7 mM cystamine, and 0.2 mM PMSF. Following incubation at 4C for 12 hr and dialysis against 10 mM Tris-HCl (pH 8.15) for 36–48 hr, the peptide/MHC monomers were purified by anion exchange (DEAE) chromatography followed by size exclusion chromatography (SEC).

Biotinylation of the purified monomers was performed using BirA enzyme. In a typical reaction, the monomer (∼40 µM) was combined with 1/10 volume of 10X BiomixA (0.5 M bicine, pH 8.3) and 1/10 volume of 10X BiomixB (100 mM ATP, 100 mM MgOAc, 500 µM d-biotin), with water added to bring the reaction to ∼400 µL. BirA was added at a ratio of 2 µg per 100 µg of substrate, and the reaction was incubated at 4C for 12 hr. Excess biotin and reaction components were removed by buffer exchange into 10 mM HEPES, 150 mM NaCl (pH 7.4) using SEC. The biotinylated NLV/A2 and GL9/A2 monomers were then incubated with their respective TotalSeq-C Streptavidin-PE (see below) at a monomer-to-streptavidin molar ratio of 5:1 to form tetramers, followed by addition of a 100-fold molar excess of biotin to block any unoccupied streptavidin sites. Finally, the tetramers were pooled together for subsequent use, as in **Fig. 3** and **Fig. S5**.

#### scTCR-seq staining, sorting, and computational pipeline

For the experiment in **Fig. 3**, cells were expanded as above for 14 days at which point a fraction of cells were first analyzed for flow cytometry using either NLV tetramer or GL9 tetramer (data shown in **Fig. 3** and **Fig. S5**). The remaining majority of cells were then prepared for single-cell analysis by counting and pooling 2-6×10^6^ live cells per condition into 5 mL FACS tubes, washing in Cell Staining Buffer (CSB, BioLegend #420201), and initially incubating in 0.04% dextran sulfate and 0.3% True-Stain Monocyte Blocker (BioLegend # 426103) for 15 min at 4C. Cells were then stained with a mixture of four barcoded tetramers (prepared as indicated above) - two distinct barcodes for NLVPMVATV (BioLegend #405297, #405295) and two distinct barcodes for GILGFVFTL (BioLegend #405161, #405163) - at a final concentration of 1 nM per tetramer in CSB for 30 min at 4C in the dark. After this time, cells were washed 3x in CSB and subsequently incubated with Human TruStain FcX Fc Receptor Blocking Solution (BioLegend #422302) and LIVE/DEAD Fixable Violet Dead Cell Stain Kit (ThermoFisher Scientific # L34964) in 1X PBS for 10 min at 4C in the dark. After this incubation, a cocktail of CD8a (BioLegend #344753) and CD4 (BioLegend #344651) CITE-seq antibodies were added to each tube and each condition was individually hashed using 1 µL of TotalSeq-C Hashtag antibody (BioLegend #394661-394671) per 100 µL stain, as outlined in **Fig. S5**, and incubated for 15 min at 4C in the dark. After this stain, cells were washed 4x with chilled CSB and transferred to clean, barcode-naïve FACS tubes after the second wash. After the last wash, cells were brought up in 500 µL CSB and sorted according to the gating scheme in **Fig. S5** on a Sony SH800 flow cytometer into collection tubes containing 1X PBS + 0.04% BSA prior to counting live cells and loading onto a Chromium X instrument (10X Genomics) using Chromium Next GEM Single Cell 5’ Kit v2 encapsulation. Gene expression and feature barcoding libraries were generated according to kit instructions and pools across conditions were sequenced on a NovaSeq6000 (Illumina).

Raw sequencing reads were aligned, a cell-feature matrix was constructed, VDJ contigs were assembled, and cells were assigned clonotypes using the 10X Genomics CellRanger software. All downstream analysis using the resulting cell-feature matrices was performed with the Seurat package (v4.3.0.1) (*60*). In brief, donor of origin was assigned for each cell on the basis of antibody hashtag reads, using the HTODemux algorithm, as described previously (*61*) and implemented in Seurat. Cells were further classified as CD4 or CD8 on the basis of anti-CD4 and anti-CD8 CITE-seq antibody reads. Finally, barcoded tetramer reads were used to assign antigen specificity to each clonotype.

### *In vitro* cytotoxicity assay

For the experiment described in **Fig. S7**, 1×10^4^ A375-mScarlet^NLV^ or A375-mScarlet^ELA^ tumor cells were plated in clear, flat-bottom 96-well plates 24 hours prior to co-culture to allow for adequate cell adherence to plastic. NLV+ CD8+ T cells were added at E:T ratios ranging from 1:1 to 1:8, where “effector” refers to NLV-specific CD8+ T cells (antigen-specific cell % determined by flow cytometry, per condition, prior to each experiment). Tumor cell growth was tracked using a live cell imager (Incucyte S3) every 3 hr for 7 days. Tumor cell killing was determined based on red object area (normalized to t=0 hr), and experiments were conducted in complete RPMI-1640 media without IL-2.

### TIL culture

Patient-derived TIL were generated from ovarian tumors (either primary or omental metastases) at time of surgical staging. For the experiments described in **Fig. 5** and **Fig. S9**, tumors were sectioned into at least 24, 2×2 mm fragments and placed in individual wells of a 48-well plate containing AIMV/RPMI media supplemented with 6000 U/mL IL-2 and left undisturbed for 2 days to allow for TIL outgrowth. After 48 hours, baseline 4-1BB expression as well as percentage of CD3, CD4, and CD8-positive TIL were measured and select wells were transduced with titrated volumes of 4-1BB ZsGreen or Fas-4-1BB-ZsGreen LV. Transduction efficiency based on ZsGreen expression was measured on days 2, 5, and 9 post-transduction.

### Tumor associated lymphocyte experiment

Patient-derived tumor associated lymphocytes (TAL) and tumor cells were generated from the ascites of patients with advanced ovarian cancer. For the experiment described in **Fig. 5**, cells were isolated via Ficoll separation and cryopreserved until ready for use. Lymphocytes and tumor cells were later thawed and placed in 48-well plates in AIMV/RPMI media supplemented with 6000 U/mL IL-2 for 48 hours, after which time the percentage of CD3, CD4, and CD8-positive cells were measured. TAL were pre-activated with anti-CD3/anti-CD28 stimulation prior to transduction with titrated volumes of 4-1BB ZsGreen LV. Transduction efficiency is based on ZsGreen expression measured on day 7 post-transduction.

### AIMV/RPMI Media for TIL culture

50% AIMV (Gibco #12055091)

50% RPMI (Gibco #11875085)

10% Human serum (Bio IVT HS1017HI)

1% Glutamax (Gibco #35050061)

12.5 mmol/L HEPES (Gibco #15630080)

1% Pen/Strep (Gibco #15140122)

5 µg/mL gentamicin (Gibco #15750060)

6000 U/mL IL-2 (Biotechne #BT-002-500)

### Flow cytometry, tetramers/dextramers, and antibodies

Cells were analyzed on a Cytoflex S (Beckman) or sorted on a Sony SH800 flow cytometer.

NLV tetramer was made in-house (described above) and coupled to APC (BioLegend #405207) or PE (BioLegend #405203) fluorophores. NLV dextramer was purchased from Immudex (cat #WB02132). U-load dextramer was used for experiments involving NLV, GLC, and GL9 (**Fig. 2**), and loaded with individual peptides purchased from GenScript (crude scale) following Immudex manufacturer’s instructions. CountBright Absolute Counting Beads (ThermoFisher Scientific # C36950) were used to determine absolute cell counts in flow cytometric assays.

For all experiments, unless otherwise stated, cells were initially transferred to 96-well V-bottom plates and first washed with 1X PBS prior to staining with 1:500 of LIVE/DEAD Fixable Viability Dye (ThermoFisher, see below for cat #s) for 10 min at 4C in the dark. All subsequent stains and washes were conducted in chilled FACS buffer (1X PBS + 0.5% bovine serum albumin + 2 mM EDTA). If cells were being assessed for antigen-specific T cell analysis, tetramer was then added to cells at 1:200 and allowed to stain for 30 minutes at 4C in the dark. If dextramer was being used, Immudex dextramer was added at 1:200 or U-load dextramer was added at 1:100, and cells were allowed to stain for 20-30 minutes at room temperature in the dark, per manufacturer’s instructions. Cells were then spun down at 300 *xg* for 5 minutes, and stained with indicated antibody cocktails containing antibodies obtained from BioLegend (see below for specific cat #s, except for TotalSeq-C antibodies which are listed above) all at 1:200 dilution from the stock solution in chilled FACS buffer and allowed to stain for 15 minutes at 4C in the dark. Cells were then washed 3x in chilled FACS buffer prior to flow cytometric analysis. Antibodies used in this manuscript:

anti-human CD3 (BioLegend, Alexa Fluor 647 #344826; BioLegend, Brilliant Violet 711 #300464;

BioLegend, Alexa Fluor 700 #317340)

anti-human CD4 (BioLegend, Pacific Blue #980806; BioLegend, APC #357408)

anti-human CD8 (BioLegend, PE #344706; BioLegend, PE/Cyanine7 #344750)

anti-human CD16 (BioLegend, PE #980102)

anti-human CD19 (BD Biosciences, BUV496 #612939)

anti-human CD25 (BioLegend, PE #302606)

anti-human CD39 (BioLegend, APC/Cyanine7 #328226)

anti-human CD56 (BioLegend, APC #981204)

anti-human CD103 (BioLegend, Brilliant Violet 421 #350214)

anti-human CD137 (4-1BB) (BioLegend, PE/Cyanine7 #309818; BioLegend, APC #309810; BioLegend, PE #309804)

anti-murine CD137 (4-1BB) (BioLegend, PE #106106)

anti-human EGFR (BioLegend, APC #352906)

CellTrace Violet Cell Proliferation Kit (ThermoFisher Scientific #C34557)

Fixable Viability Kit (BioLegend, Zombie Yellow #423104)

LIVE/DEAD Fixable Violet Dead Cell Stain Kit (ThermoFisher Scientific #L34964)

LIVE/DEAD Fixable Yellow Dead Cell Stain Kit (ThermoFisher Scientific #L34959)

### Software and statistical analysis

Graphs presented in this manuscript were generated using GraphPad Prism 10 and data were analyzed for statistical significance using built-in GraphPad Prism software with the appropriate tests specified in the corresponding figure legends. Results were considered significant if *P* < 0.05. Flow cytometry data were analyzed using FlowJo (v10.10.0).

## List of Supplementary Materials

**Fig. S1. – Fig. S9** – supplementary figures referenced throughout the manuscript.

**Table S1** – 4-1BB targeting plasmid sequences used in this study (pMD2_type I, type II, scFvs).

**Table S2** – Transfer/cargo plasmid sequences used in this study (pBS-ZsGreen, pHIV-mScarlet).

**Table S3** – MHC class I peptides used in this study (95 CEF).

## Acknowledgments

For technical support, we thank the Koch Institute’s Robert A. Swanson (1969) Biotechnology Center, particularly the Flow Cytometry Facility and MIT BioMicro Center for their support. We also thank Catherine J. Wu, Eric Smith, Kai Wucherpfennig, Max Jan, Stefani Spranger, and Arlene Sharpe for their helpful discussions throughout this work. Schematic figures in the manuscript were created with BioRender.com.

