## Supplementary Materials for "Activation-dependent lentiviruses enable antigen-specific T cell expansion and transduction"

### Supplementary Figures

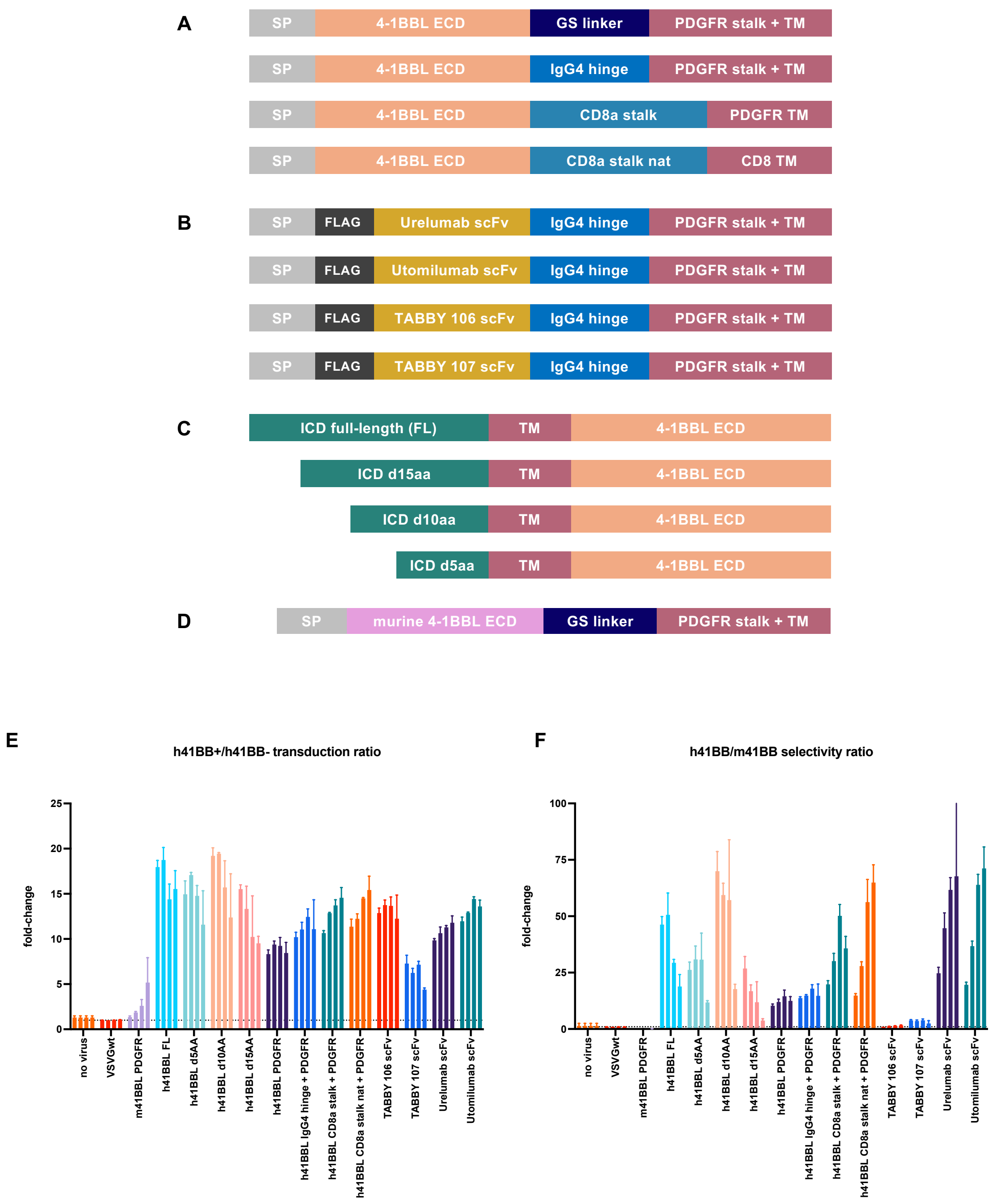

**Fig S1.** **(A)** Architectures of 4-1BB-targeting constructs to be displayed on the surface of a LV particle**.** (A) Type I TM 4-1BBL series. SP = signal peptide (IGHM for all constructs, see **Table S1** for details). **(B)** Type I TM formatted scFv series. **(C)** Type II TM 4-1BBL series. **(D)** murine 4-1BBL (m4-1BBL) formatted as a type I TM protein. (**E**) Briefly, h41BB+/h41BB- transduction ratio is calculated as the on-target (4-1BB+) transduction % / off-target (4-1BB-) transduction %. Each virus was added in a dilution series with dextran, with each bar from left to right (per construct) representing a virus dose of: 1, 0.25, 0.0625, and 0.015625 µL from left to right as in Fig. 1. See Materials and Methods for more. (**F**) h41BB/m41BB selectivity ratio is calculated as the % h4-1BB+ ZsG+ / % m4-1BB+ ZsG+ per matched virus dose (1, 0.25, 0.0625, and 0.015625 µL).

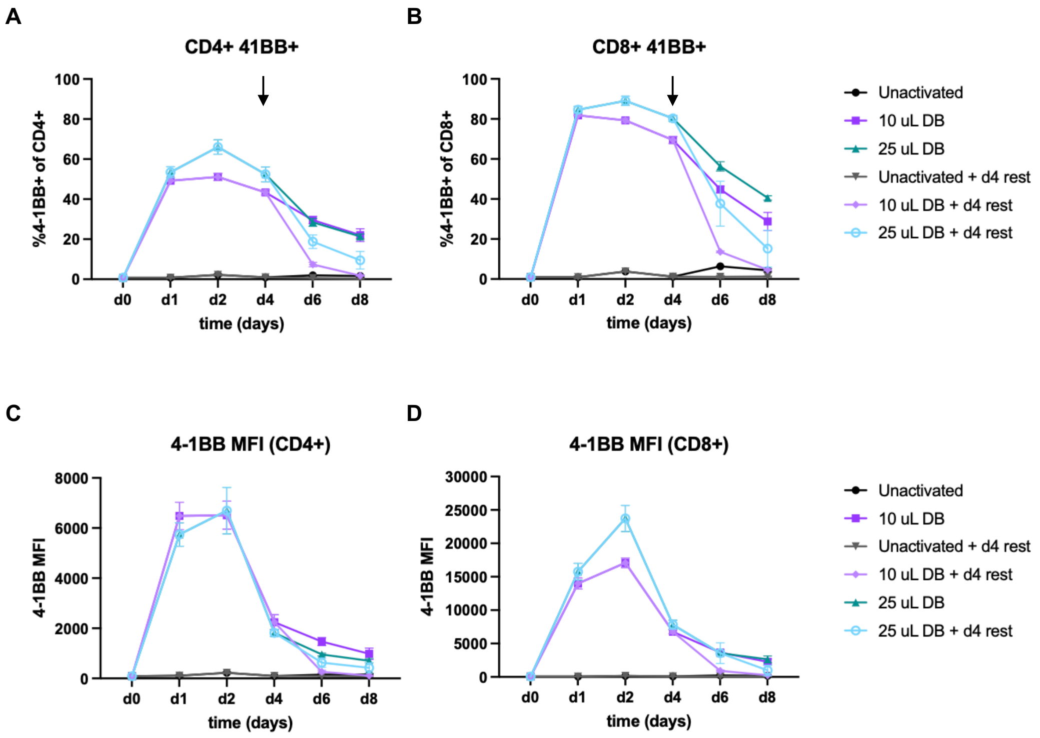


**Fig S2.** **(A)** Line plots of 4-1BB receptor expression in CD4+ T cells and **(B)** in CD8+ T cells by percentage, over time. **(C)** Line plots of 4-1BB receptor expression in CD4+ T cells and **(D)** in CD8+ T cells by mean fluorescent intensity (MFI), over time. Arrows in **(A)** and **(B)** indicate the timepoint (96 hr, d4) at which point anti-CD3/anti-CD28 Dynabead (DB) stimulation was withdrawn in the "+d4 rest" conditions. Otherwise, "DB" conditions maintained continuous anti-CD3/anti-CD28 stimulation (either 10 µL or 25 µL of DBs) throughout the timepoints assessed.

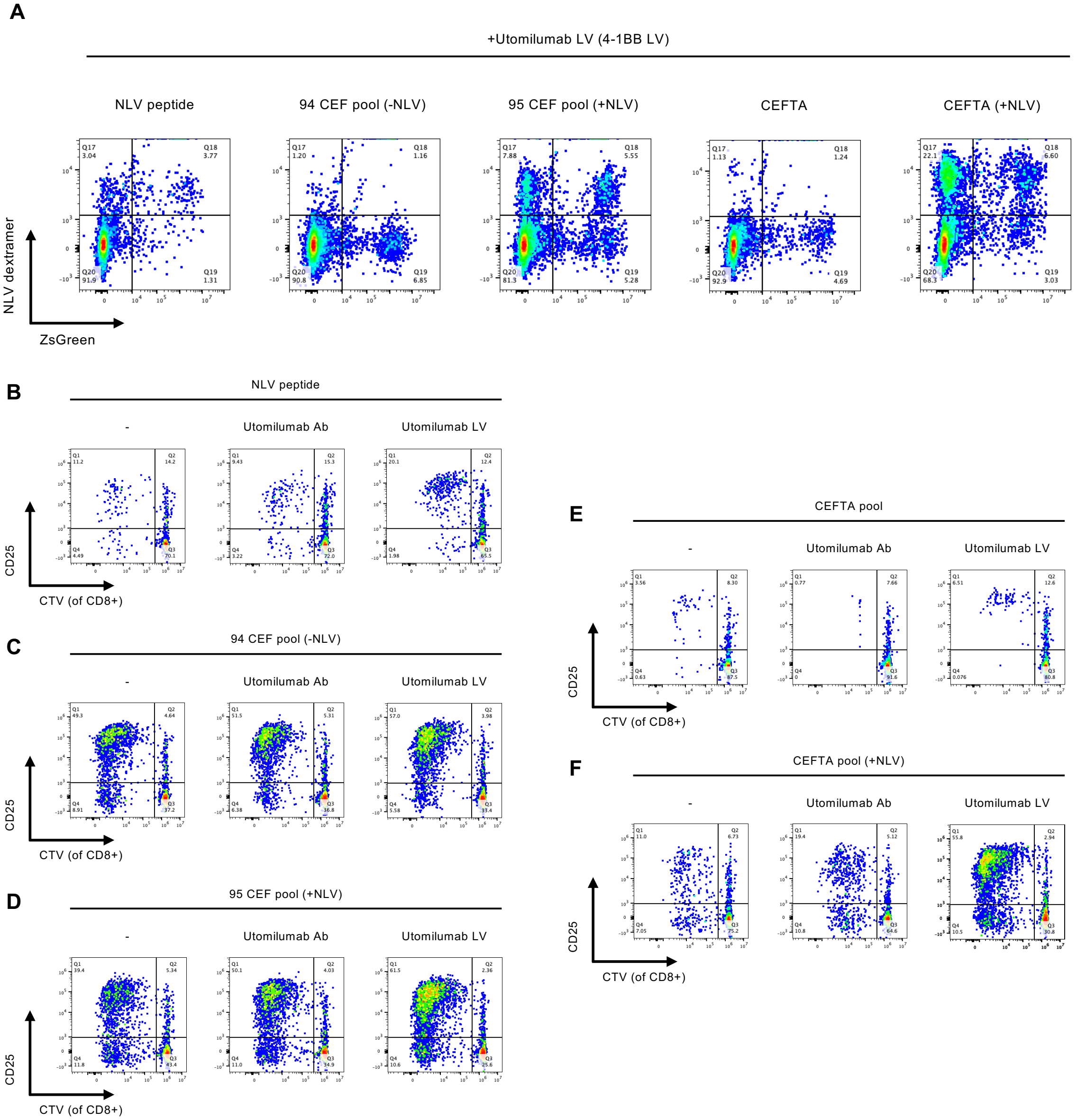


**Fig S3. (A)** Representative flow plots of PBMCs grown in the presence of NLV peptide only, MHC class I 94-member CEF library lacking NLV peptide (-NLV), MHC class I 95-member CEF library containing NLV peptide (+NLV), MHC class II CEFTA library lacking NLV peptide, or MHC class II CEFTA library containing NLV peptide (+NLV). After 24 hours of stimulation, 4-1BB LV (Utomilumab LV) was added to the indicated conditions and transduction of antigen-specific T cells was measured by NLV dextramer+ ZsGreen+ expression after 7 days. **(B)** Representative flow plots of PBMCs grown in the presence of NLV peptide only (-), Utomilumab antibody (1 μg/mL) or Utomilumab LV (MOI = 2), assessing activation (CD25+) and proliferation (CTV low) in CD8+ T cells (gated on live+, CD8+), in the absence of dextran. CTV = CellTrace Violet. **(C)** Same as in (B), however PBMCs were grown in the presence of a 94-member CEF library lacking NLV peptide. **(D)** Same as in (B) & (C), however PBMCs were grown in the presence of a 95-member

CEF library containing NLV peptide. **(E)** Same as in (B), however PBMCs were grown in the presence of a 35-member CEFTA library lacking NLV peptide. **(F)** Same as in (E), however PBMCs were grown in the presence of a 35-member CEFTA in addition to NLV peptide.

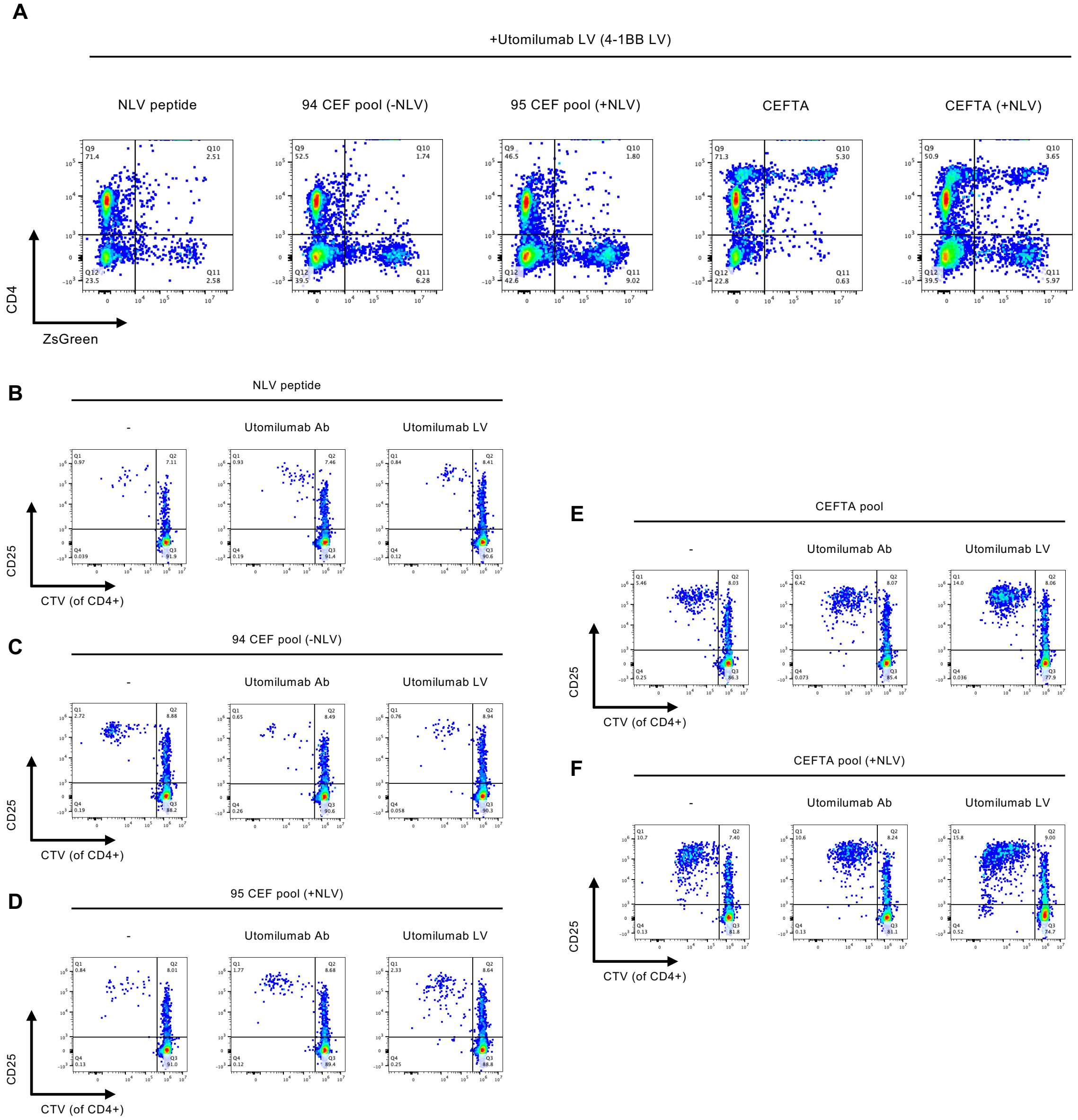


**Fig S4. (A)** Representative flow plots of PBMCs grown in the presence of Utomilumab LV (4-1BB LV) after stimulation with the indicated peptide(s), assessing transduction (ZsGreen+) in CD4+ T cells (gated on live+), in the absence of dextran, after 7 days. **(B)** Representative flow plots of PBMCs grown in the presence of NLV peptide only (-), Utomilumab antibody (1 µg/mL) or Utomilumab LV (MOI = 2), assessing activation (CD25+) and proliferation (CTV low) in CD4+ T cells (gated on live+, CD4+), in the absence of dextran. **(C)** Same as in (B), however PBMCs were grown in the presence of a 94-member CEF library lacking NLV peptide. **(D)** Same as in (B) & (C), however PBMCs were grown in the presence of a 95-member CEF library containing NLV peptide. **(E)** Same as in (B), however PBMCs were grown in the presence of a 35-member CEFTA library lacking NLV peptide. **(F)** Same as in (E), however PBMCs were grown in the presence of a 35-member CEFTA, alongside NLV peptide.

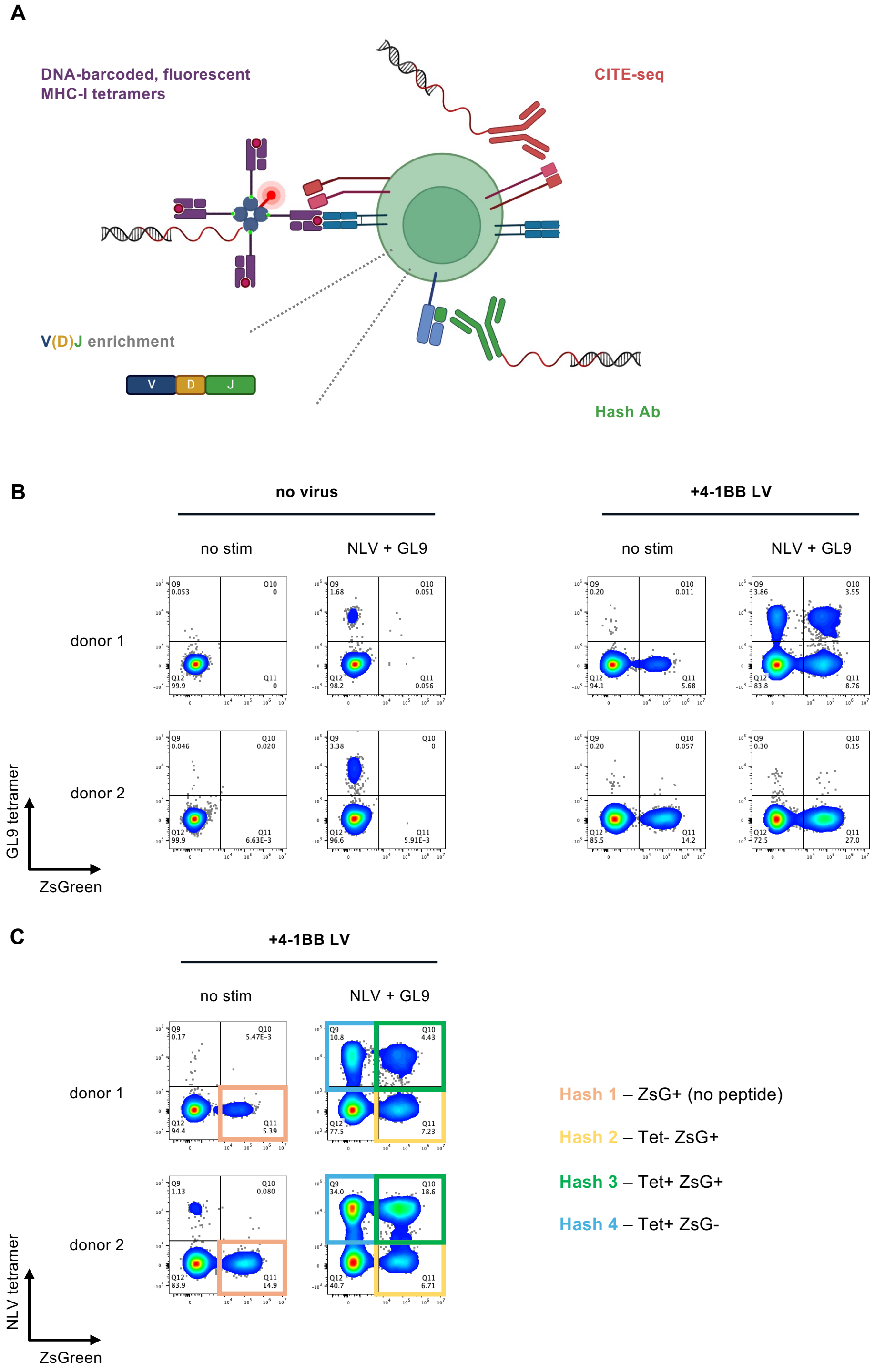

**Fig S5. (A)** Schematic of scTCR-seq workflow. Individual T cells are defined by lineage using anti-CD4 or anti-CD8a CITE-seq antibodies, antigen specificity via DNA-barcoded MHC I tetramers, clonotype through V(D)J enrichment, and can be pooled across conditions using hashing antibodies. **(B)** Representative flow plots of PBMCs grown in the absence (no stim) or presence of indicated peptides (NLV + GL9 peptide pool), in the absence (no virus) or presence of 4-1BB LV with dextran. Conditions were assessed for GL9-specific expansion (GL9 tetramer+) and transduction (ZsGreen+) prior to scTCR-seq, gated on live cells, at day 14 in donor 1 (top row) and donor 2 (bottom row). **(C)** Sorting scheme for scTCR-seq experiment in **Fig. 3**. Four different conditions were individually hashed (as indicated) and sorted prior to pooling cells for scTCR-seq.

**A**

| Donor 1 clonotypes | CDR3b | CDR3a | ZsG+ (no peptide) | Tet+ ZsG+ | Tet+ ZsG- | Tet- ZsG+ | Antigen specificity | VDJdb |
|---|---|---|---|---|---|---|---|---|
| clonotype1 | CASSLVAGVPNEQFF | CAVRYGNDMRF | 1.3 | 23.8 | 34.6 | 1.0 | NLV | |
| clonotype3 | CSARVEFASGSPAIYNEQFF | CAMSAEDDKIIF | 0.2 | 13.1 | 5.5 | 10.8 | NLV | |
| clonotype2 | CASSIRSSYEQYF | CAAGGSQGNLIF | 0.3 | 12.9 | 9.0 | 5.9 | GL9 | GL9 |
| clonotype11 | CASSQACRTDTQYF | | 27.9 | 0.0 | 0.0 | 0.0 | | |
| clonotype4 | CASSYSTGATFNYGYTF | CAYNAGNMLTF | 0.2 | 2.6 | 7.3 | 2.4 | NLV | NLV |
| clonotype5 | CASSPGLGQMFGETQYF | CATIHSGYSTLTF | 0.2 | 1.3 | 9.3 | 0.3 | NLV | |
| clonotype6 | CASSQVTGTPSEKLFF | CAVGFGNQFYF | 0.2 | 3.5 | 5.5 | 1.9 | NLV | |
| clonotype8 | CASSVLETGFDEQFF | CAVPPGGNNDMRF | 0.0 | 3.2 | 3.8 | 3.7 | | |
| clonotype10 | CASSQLGGAGLINEQFF | CAVKNFNKFYF | 0.0 | 3.5 | 2.7 | 3.8 | | |
| clonotype7 | CASSLWDRGSGANVLTF | CAARSNFGNEKLTF | 0.5 | 3.6 | 5.6 | 0.0 | NLV | |

**B**

| Donor 2 clonotypes | CDR3b | CDR3a | ZsG+ (no peptide) | Tet+ ZsG+ | Tet+ ZsG- | Tet- ZsG+ | Antigen specificity | VDJdb |
|---|---|---|---|---|---|---|---|---|
| clonotype1 | CASSPTSGSPGELFF | CAVFYGNKLVF | 0.2 | 89.3 | 89.9 | 31.0 | NLV | |
| clonotype2 | CASSQEGSGANVLTF | CAFLPLMYSGGGADGLTF | 13.1 | 0.0 | 0.0 | 15.4 | | |
| clonotype3 | CASSYAWGLNHTEAFF | CAVEGTYKYIF | 7.1 | 0.4 | 0.0 | 5.0 | | |
| clonotype10 | CSARSLDAGSSYNEQFF | CAVDNYGQNFVF | 7.0 | 0.0 | 0.0 | 0.0 | | KLG |
| clonotype8 | CASADRAFGYTF | CAMSGVYSGAGSYQLTF | 0.5 | 0.0 | 0.0 | 6.0 | | |
| clonotype18 | CASTIEGLAPYEQYF | CAFMKLMYSGGGADGLTF | 4.2 | 0.4 | 0.0 | 0.0 | | |
| clonotype11 | CASSLEGGESYEQYF | CAVGAVKYSGGGADGLTF | 2.4 | 0.0 | 0.0 | 1.1 | | |
| clonotype13 | CASSPEGGSGNTIYF | CAETKVMYSGGGADGLTF | 1.9 | 0.0 | 0.0 | 1.7 | | |
| clonotype4 | CASGGPFGTDTQYF | CAFITGNQFYF, CAVRYNNDMRF | 0.0 | 2.1 | 3.5 | 1.5 | NLV | |
| clonotype15 | CSAGLAGRNNEQFF | CAVEPSGSARQLTF | 2.9 | 0.0 | 0.0 | 0.3 | | |

**Fig S6. (A)** CDR3b and CDR3a sequences of the top 10 expanded clonotypes in donor 1 and **(B)** donor 2 from the experiment in **Fig. 3**. The proportion of each clonotype (row) sequenced per condition (hash, column) is indicated as a percentage. Antigen specificity of each clonotype is listed (if detected), and clonotypes were passed through VDJdb (*42*) to assess for newfound antigenic specificity. VDJdb can be accessed at: https://vdjdb.cdr3.net/.

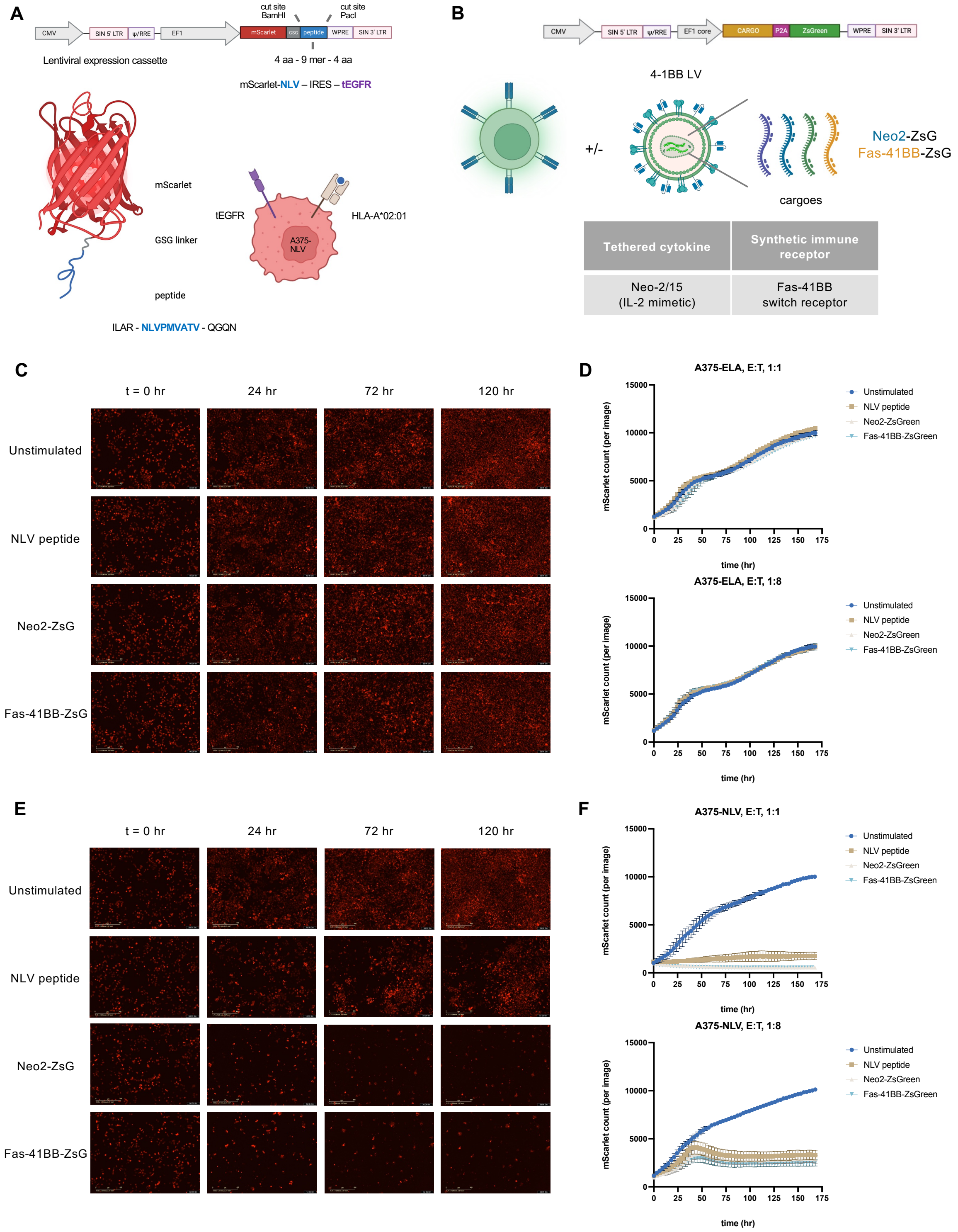


**Fig S7. (A)** A lentiviral cassette was designed for stable expression of mScarlet-antigen in tumor cells. The MHC class I peptide of interest (in blue) is flanked by ~4 amino acid residues of native sequence on each side of the epitope to promote canonical antigen processing (e.g. the

NLVPMVATV antigen of interest is flanked with native N-terminal residues “ILAR” and C-terminal residues “QGQN”). Restriction enzyme cut sites for BamHI and PacI have been placed flanking the peptide of interest to leave mScarlet scarless, by embedding BamHI within the flexible GSG linker and PacI immediately following the stop codon of the chimeric protein for facile cloning of epitopes of interest. The lentiviral cassette also possesses a truncated EGFR (tEGFR) marker protein, following an IRES sequence, for orthogonal detection of mScarlet-transduced tumor cells in the event of antigen loss. A375 melanoma cells are HLA-A*02:01-expressing tumor cells. Transduction with mScarlet-NLV antigen, results in an antigen-presenting A375-mScarlet$^{NLV}$ tumor line. **(B)** The designed lentiviral cassette places a genetic cargo of interest (listed in the table below) in front of a self-cleavable P2A sequence and a ZsGreen fluorophore for facile detection of transduced cells. **(C)** “Off-target” A375-mScarlet$^{ELA}$ tumor cells are co-cultured with PBMCs that were expanded with no peptide (unstimulated), NLV peptide only (NLV peptide), or NLV peptide + anti-4-1BB LV containing engineered cargoes (Neo2-ZsG or Fas-41BB-ZsG). mScarlet tumor cells are visualized over the course of 7 days (168 hr) and tumor cell growth is tracked and analyzed by an Incucyte S3 live cell imager. Representative images represent the indicated cells co-cultured with A375-mScarlet$^{ELA}$ tumor cells at an effector-to-target (E:T) ratio of 1:1. **(D)** mScarlet tumor growth curves plotted at an E:T of 1:1 and 1:8 in A375-mScarlet$^{ELA}$ tumor cells. **(E)** and **(F)** same as in (C) and (D), respectively, but with “on-target” A375-mScarlet$^{NLV}$ tumor cells.

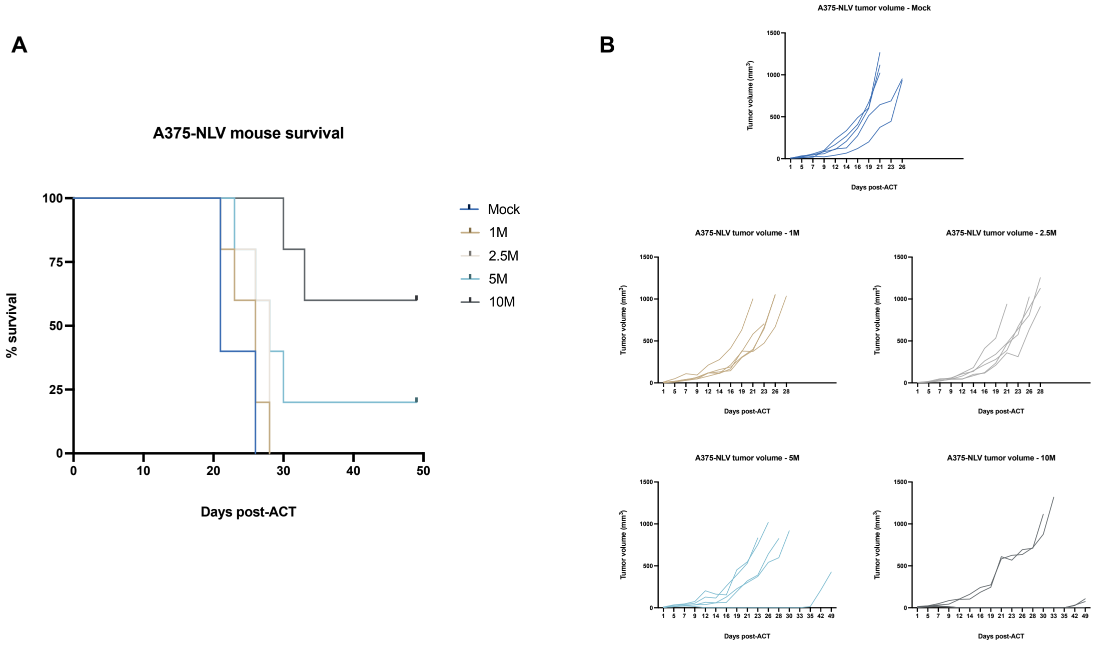


**Fig S8. (A)** A375-mScarlet$^{NLV}$ tumor-bearing mice received no T cells (mock), or $1x10^6$, $2.5x10^6$, $5x10^6$, or $10x10^6$ peptide-expanded NLV+ CD8+ effector T cells in a dose-finding pilot study. Kaplan-Meier curve for overall survival. N=5 mice per tested T cell dose. **(B)** Individual tumor curves for each experimental cohort in which 1 mouse in the $5x10^6$ and 3 mice in the $10x10^6$ cohorts achieved a complete response during the study.

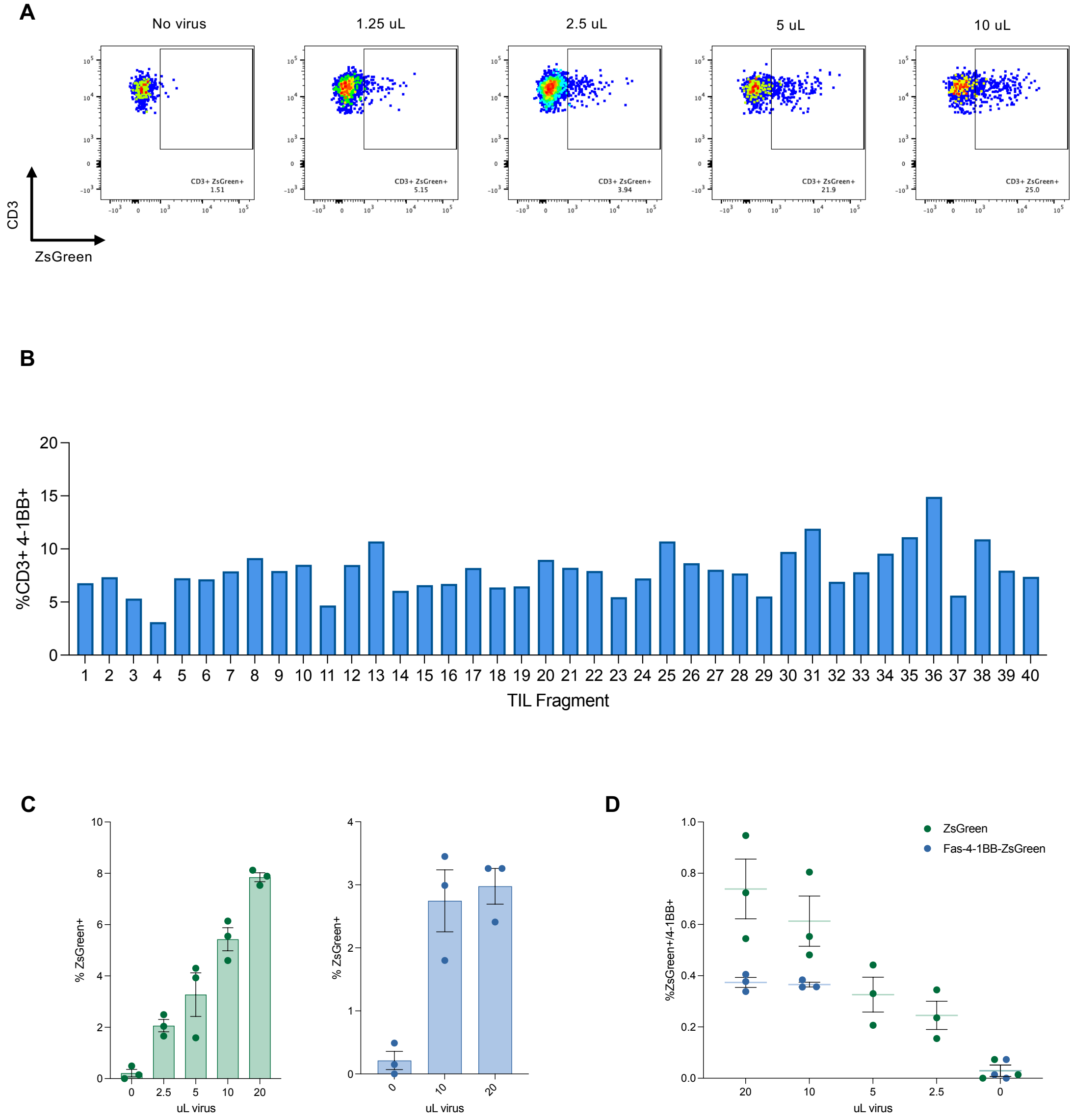


**Fig S9. (A)** Representative flow plots for the data presented in **Fig. 5D-E**. **(B)** Bar plots summarizing the percentage of CD3+, 41BB+ T cells from 40 different tumor fragments derived from a metastatic lesion resected from the omentum of a patient with ovarian cancer after 2 days in culture. **(C)** Left: 4-1BB LV containing ZsGreen was added to triplicate tumor fragments at four different doses (20, 10, 5, or 2.5 μL) and transduction (%ZsGreen+) was compared to no virus tumor fragments. Right: 4-1BB LV containing Fas-4-1BB-ZsGreen was added to triplicate tumor fragments at two different doses (20 or 10 μL) and transduction (%ZsGreen+) was compared to no virus tumor fragments. **(D)** Transduced TILs (ZsG$^{+}$) at day 2 are compared to their matched, baseline 4-1BB+ expression at day 0, representing the fraction of 'transducible' TIL (%ZsGreen/4-1BB+) achieved across a range of LV doses and two different 4-1BB LVs (ZsGreen or Fas-4-1BB ZsGreen).
